# Unraveling single-molecule reactions via multiplexed in-situ DNA sequencing

**DOI:** 10.1101/2025.11.27.690701

**Authors:** Jagadish Prasad Hazra, Rebecca Andrews, Piers Turner, Qing Zhao, Horst Steuer, Afaf H. El-Sagheer, Abhishek Mazumder, Hafez El Sayyed, Mirjam Kümmerlin, Tom Brown, Achillefs N. Kapanidis

## Abstract

DNA sequence regulates complex reactions and protein-DNA interactions, yet sequence effects remain poorly understood due to the lack of direct, high-throughput approaches to study sequence-dependence at the single-molecule level. Here, we introduce Single-molecule Phenotyping and In-Situ Sequencing (SPIN-Seq), a DNA-based, protein-free method that links functional and structural properties of a single DNA molecule with the sequence of that same molecule. After performing functional assays on immobilized DNA molecules, SPIN-Seq uses sequencing-by-transient-hybridization on the same surface and instrument to read sequences in each immobilized DNA molecule, directly linking phenotype and genotype. Applying SPIN-Seq, we dissected the interaction of a transcription factor with its target sequence, and revealed the sequence-dependence of RNA polymerase pausing and reaction-path branching during initial transcription. Our method provides powerful, systematic ways to understand complex molecular mechanisms and their sequence dependence.

---

Interactions of nucleic acids with proteins and other biomolecules are key for genetic processes and biotechnology. Analysis of such interactions relies heavily on structural, biochemical, and biophysical approaches; single-molecule methods, in particular, offer key insight on nucleic acid interactions, often capturing them in real-time (*1–6*). However, single-molecule experiments typically study one sequence at a time and do not offer a global view of how DNA sequence affects protein-DNA interactions.

An attractive route to explore sequence-dependence at single-molecule level couples single-molecule DNA sequencing (nanopore- or fluorescence-based) to analysis of protein-driven processes on DNA strands being sequenced (*7–9*); however, these approaches are confined to proteins translocating on nucleic acids, and some require special surfaces. Recently, two approaches enabled multiplexing on DNA-DNA and protein-DNA interactions (*10, 11*); both involved single-molecule fluorescence imaging on custom-built microscopes, DNA amplification, Illumina sequencing, and linking of single-molecule and sequence readouts. While these powerful approaches offered insight on how sequence controls the structure and dynamics of folded DNA and protein-DNA complexes, they had not addressed labelled proteins or non-equilibrium reactions and their kinetics, in part due to their dependence on a solid support (MiSeq chips) suboptimal for single-molecule imaging.

Here, we introduce Single-molecule Phenotyping and In-Situ Sequencing (SPIN-Seq; **Fig. 1**), a method that links the functional properties of a single DNA molecule (its “single-molecule phenotype”) with the sequence of the *same* molecule, on the same surface and instrument. This phenotype involves structure and dynamics, e.g., distances within protein-DNA complexes, or kinetics of reactions and interactions. SPIN-Seq starts by immobilizing DNA libraries on standard glass coverslips; DNAs then interact with proteins, and dynamics are recorded; after protein removal, a new sequencing-by-hybridization approach reads sequences in each DNA molecule, and links them to its single-molecule phenotype to gain mechanistic insight. Our assay can read up to 5 nucleotides in length, covering libraries of up to 1024 DNA sequences and opening a new avenue for exploring protein-DNA interactions.

**Figure 1.**
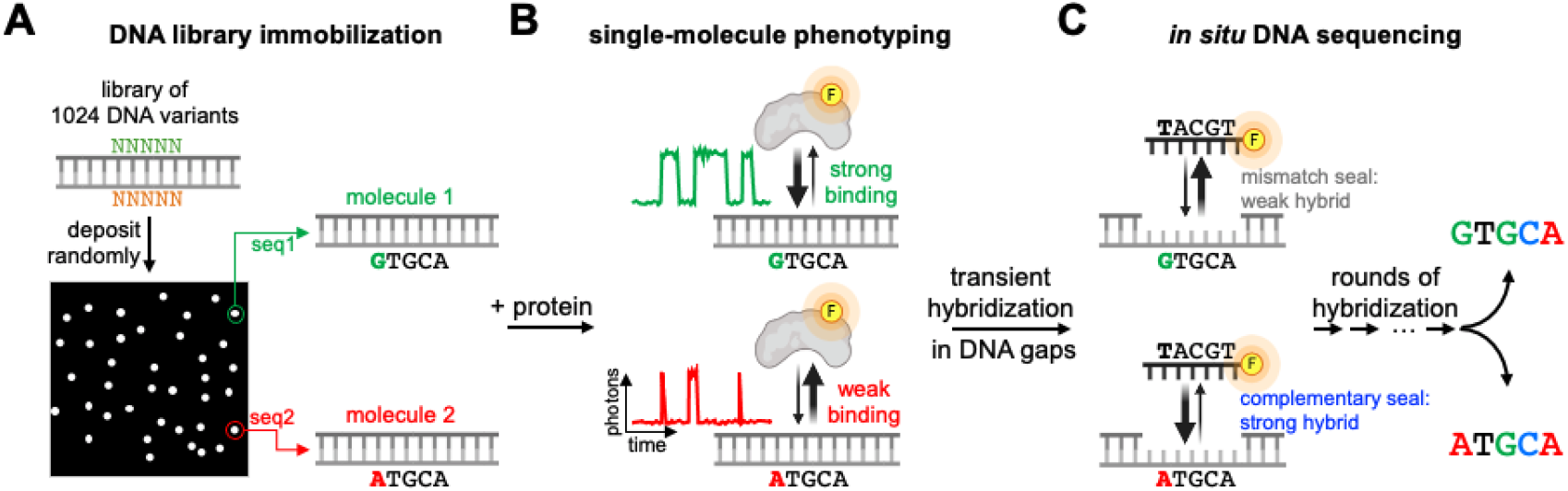
A single-molecule method that links protein-DNA interactions to DNA sequence. **(A)** A library of DNA molecules (e.g., with 4^5^ DNA sequences, reflecting a 5-nt randomized sequence NNNNN) is immobilized randomly on a surface, and imaged using widefield single-molecule fluorescence microscopy. Different molecules (white filled circles) contain different DNA sequences. **(B)** Proteins are added, and the single-molecule kinetics of protein-DNA interactions (its “single-molecule phenotype”) are recorded using fluorescence signals (here, due to binding of labelled proteins). **(C)** Surface-immobilized DNA molecules are converted into gapped DNA molecules *in situ*. Short oligonucleotides (DNA “seals”) are added and bind transiently to the gap; the extent of binding differs between fully complementary seals and mismatched seals, forming the basis for recovering the sequence of each molecule and linking it to single-molecule phenotypes.

## Single-molecule hybridization on DNA gaps

To perform *in situ* single-molecule sequencing, we leveraged the transient binding of fluorescent single-stranded (ss) DNA oligonucleotides (“seals”) to surface-immobilized DNAs featuring short ssDNA gaps (*12*) (**Fig. 1**, right; **Fig. 2A-B**). Such transient binding is used in high-throughput endeavors, such as DNA-PAINT (*13*). We selected gapped-DNA templates instead of ssDNA, since the coaxial stacking at the gap ends stabilizes the bound seal; the gap also defines the sequenced segment. Base-calling relies on distinguishing which DNA seals are fully complementary to the gap (as opposed to seals that are mismatched). The short gap length allows bound seals to thermally dissociate and be replaced by seals from the surrounding solution; repeated seal binding increases statistics and interrogation accuracy, while eliminating photobleaching.

**Figure 2.**
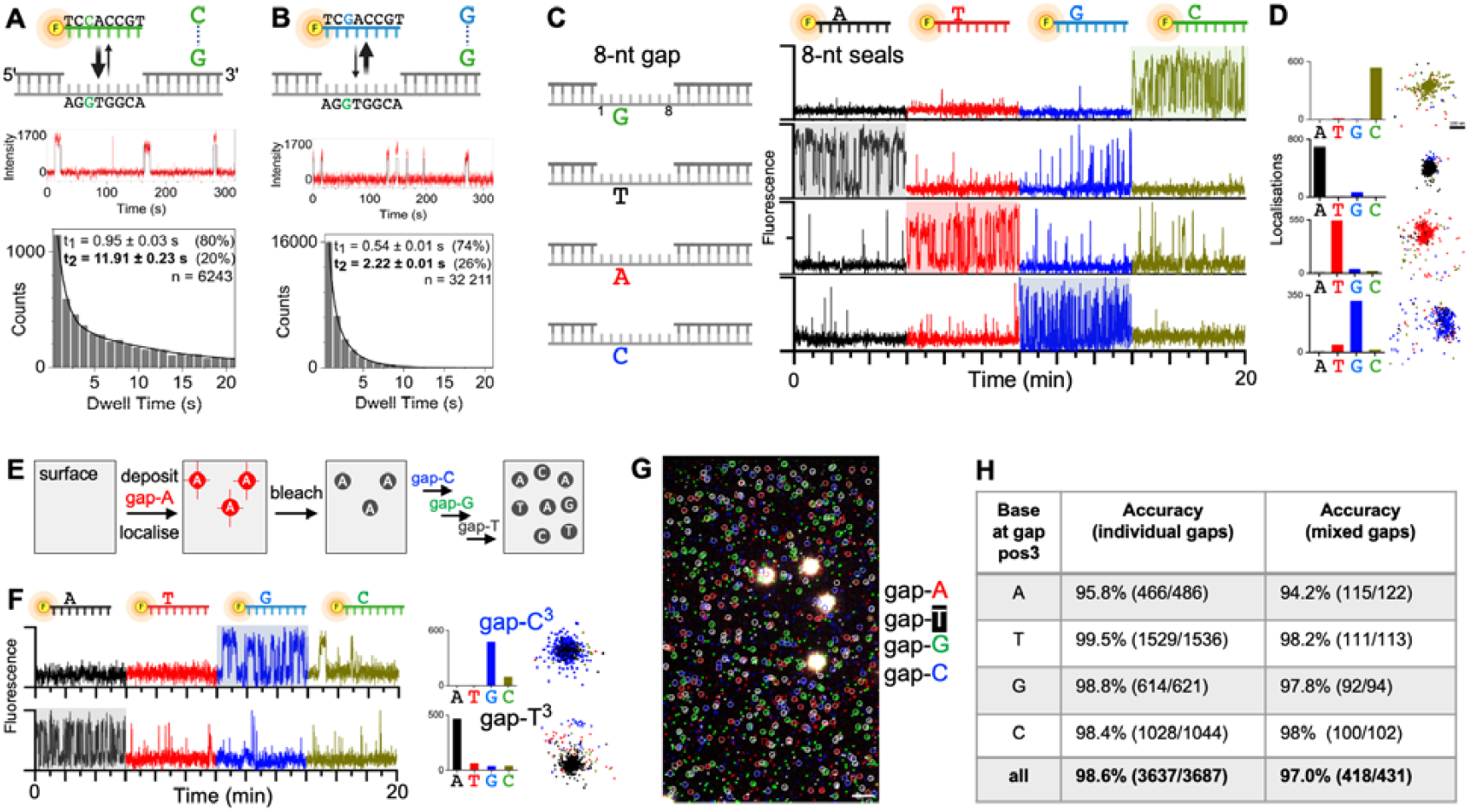
Reading single DNA bases via transient DNA binding of short DNA oligos to gapped DNA. **(A) Top**: Schematic of a fully complementary 8-nucleotide (nt) seal binding in 8-nt gapped-DNA. **Middle**: fluorescence intensity time-traces (red) for ATTO647N-labelled DNA seal binding to gapped DNA. Traces are fitted (black) using Hidden Markov Modelling (HMM), identifying bound states (high-intensity dwells) and their dwell times. **Bottom**: dwell-time distribution for bound states. Data fit best to double-exponential distributions. n: number of binding events. **(B)** As in (A), but for a G:G mismatch at gap position 3 due to the use of a different seal; the mismatch drastically destabilises gap-seal binding. **(C)** First row: an 8-nt DNA gap within an immobilised molecule is sequentially interrogated at gap position 3 (where the base is a G) using 4 seals that differ only on the base opposed to the interrogated position. Only the seal featuring a C at position 3 (green DNA) shows strong gap binding; the remaining seals show negligible binding. Rows 2-4: as in row 1, but for gaps featuring a T, A, or C at gap position 3. Only the fully complementary seal shows substantial binding. **(D)** Left: Number of single-molecule localisations per seal interrogation. Right: map of localisations clustering around the interrogated gap DNA molecule. **(E)** Analysis of surfaces with mixed sequences. Sequential deposition (using cycles of deposition, localisation, and bleaching) of 4 different DNA gap molecules of known sequence. **(F)** Sequential interrogation of the deposited DNA by seals specific for gap position 3. The extent of binding in the top and bottom traces identifies a Gap-C^3^ and a Gap-A^3^ molecule, respectively. **(G)** Field of view (50×80 μm) of molecules sequenced for gap position 3 in the experiment in panel F. Circled molecules were identified correctly; in-square molecules were assigned incorrectly. White spots: fiducial markers. Scale bar, 5 μm. **(H)** Base-calling accuracies for interrogation of single sequences (as in panel C) and of a mixture of 4 sequences (as in panel F).

To read sequence information in gapped DNA via transient seal binding, we examined the binding between a surface-immobilized 8-nt gapped DNA and labelled 8-nt seals fully complementary to the gap (**Fig. 2A; table S1** for all DNA sequences). When a seal binds to the gap, the fluorescence intensity increases to a high value lasting until a seal dissociates (**Fig. 2A**, middle). Analysis of the dwell-time distribution (**Fig. 2A**, bottom) revealed two dwell times, the longer of which signifies stable binding (t_2_ ~12 s). Experiments using an 8-nt seal with a 1-nt mismatch (G:G) to gap position 3 (**Fig. 2B**) showed >5-fold shorter binding (t_2_~2.2 s), confirming that a single-nt mismatch drastically affects seal binding, as seen for other short hybrids (*14*), and offering a basis for short-read single-molecule sequencing.

### Sequencing single bases via transient DNA hybridization

We then explored reading one base on a *single* immobilized gapped-DNA molecule. Using a modified hybridization buffer (to increase sampling of seal binding; *Methods*), we interrogated a G base at position 3 in an 8-nt gap molecule (Gap8-G^3^; superscript denotes the interrogated gap position; **Fig. 2C**, left) by sequentially incubating it with 4 seals; each seal differs only in the base opposed to the interrogated gap base. Each seal with a 1-nt mismatch (seals S8-A^3^, S8-T^3^, S8-G^3^) showed negligible or very weak gap binding (**Fig. 2C**, top row; **fig. S1**). In contrast, we observed extensive binding of the seal fully complementary to the 8-nt gap (seal S8-C^3^, forming a C:G pair at position 3; **Fig. 2C**, top row, green trace). We obtained similar results for all bases at gap position 3 (**Fig. 2C** and **fig. S1**), supporting the generality of the approach.

To characterize gap-seal interactions, we measured seal dwell times and binding frequency for all gap-seal combinations (**fig. S2**). Complementary seals bound 1.3- to 5.5-fold longer and 4- to 23-fold faster than seals with 1-nt mismatches. Overall, affinity differences between complementary and 1-nt mismatch seals ranged from 8- to 45-fold (**fig. S2**). Different mismatches disrupted binding to a different degree. Mismatches containing guanines are the least disruptive, with G:G and T:G reducing affinity only by 8- and 10-fold, respectively. In contrast, mismatches containing cytosines are the most disruptive, with C:A and C:C reducing affinity by 36- and 45-fold, respectively (**figs. S2-3**). Our results agree with ensemble measurements ((*15*); **fig. S3a**), and computational predictions of thermodynamic stability ((*16, 17*); **fig. S3b**), offering insight on canonical and non-canonical base-pairings and nucleic-acid interactions.

To “call” the interrogated base, we counted single-molecule localizations of different seals to a single gap (**Fig. 2D** and **fig. S4A**) and selected as the complementary seal the one with most localizations (**fig. S5**; *Methods*). This analysis was ~98.8% accurate (614 out of 621) in detecting a G at position 3 and similarly accurate for the remaining 3 bases (~98.6%; **Figs. 2D** and **2H**; **fig. S4B-D**).

To demonstrate our ability to address a mixed population of sequences, we sequentially deposited each single-nucleotide variant for gap position 3 on the same surface, generating the “ground truth” for the sequence of each molecule (**Fig. 2E**). We interrogated the surface as before (**Fig. 2F**), mapped read bases to the pre-registered base identities (**Fig. 2G)**, and demonstrated a ~97% base-calling accuracy (**Fig. 2H**), establishing that we can read any standard base at a single position with high accuracy.

### Sequencing single bases using competitive inhibition of gap hybridization

To sequence multiple bases, we had to account for the unknown sequence context around each interrogated position and simplify the set of seals to make the assay scalable. We thus introduced a competitive-inhibition version of our assay that uses DNAs containing degenerate bases (“N”) that feature 25% of each standard base at a single position. Each interrogation uses a mixture of two seal DNAs competing for gap binding: a reporter seal (“R-seal”; either singly labelled and fluorescent prior to binding, or fluorogenic and quenched prior to binding) that binds the gap well and contains degenerate bases at all positions examined (**Fig. 3A**, left). The second seal is unlabelled (“U-seal”); when fully complementary to the gap, it outcompetes the R-seal, suppressing its gap binding and the associated fluorescence (**Fig. 3A**, middle). However, if the U-seal contains even a single mismatch to the gap sequence, it is outcompeted by R-seal, leading to high fluorescence (**Fig. 3A**, right). For every position examined, we perform 4 sequential interrogations, each involving a mix of the R-seal with one of 4 U-seals (each U-seal containing one of the 4 standard bases at the position opposite to the interrogated site); only the complementary U-seal reduces R-seal binding, thus reading the base.

To test our approach, we examined an 8-nt gap (G8-G^3^) using an 8-nt R-seal with a degenerate base opposed to gap position 3 (R8-N^3^), and four 8-nt U-seals, each featuring one of the standard bases opposed to gap position 3 (**Fig. 3B**). Using optimized R-seal and U-seal concentrations (**figs. S6-7**), we tracked R-seal binding to the gap while it is sequentially added mixed with one of the four U-seals; we observed very low R-seal binding for the mixture containing the complementary U-seal (U8-C^3^), and substantial binding for the 1-nt mismatched seals (**Fig. 3B**, top, and **fig. S8C**). Using localization counting for base-calling (*Methods*), we detected 500 out of 502 molecules exhibiting lower R-seal binding for U8-C^3^ compared to the other U-seals, yielding a ~99.6% base-calling accuracy. Overall, we obtained high accuracy for all bases at gap position 3 (~97.1%; **Fig. 3B-D**, bottom and **fig. S8; table S2)**, as well as for mixed sequences (~98.5%; **fig. S9** and **table S2**), establishing that competitive inhibition can be used for reading single positions with high accuracy.

**Figure 3.**
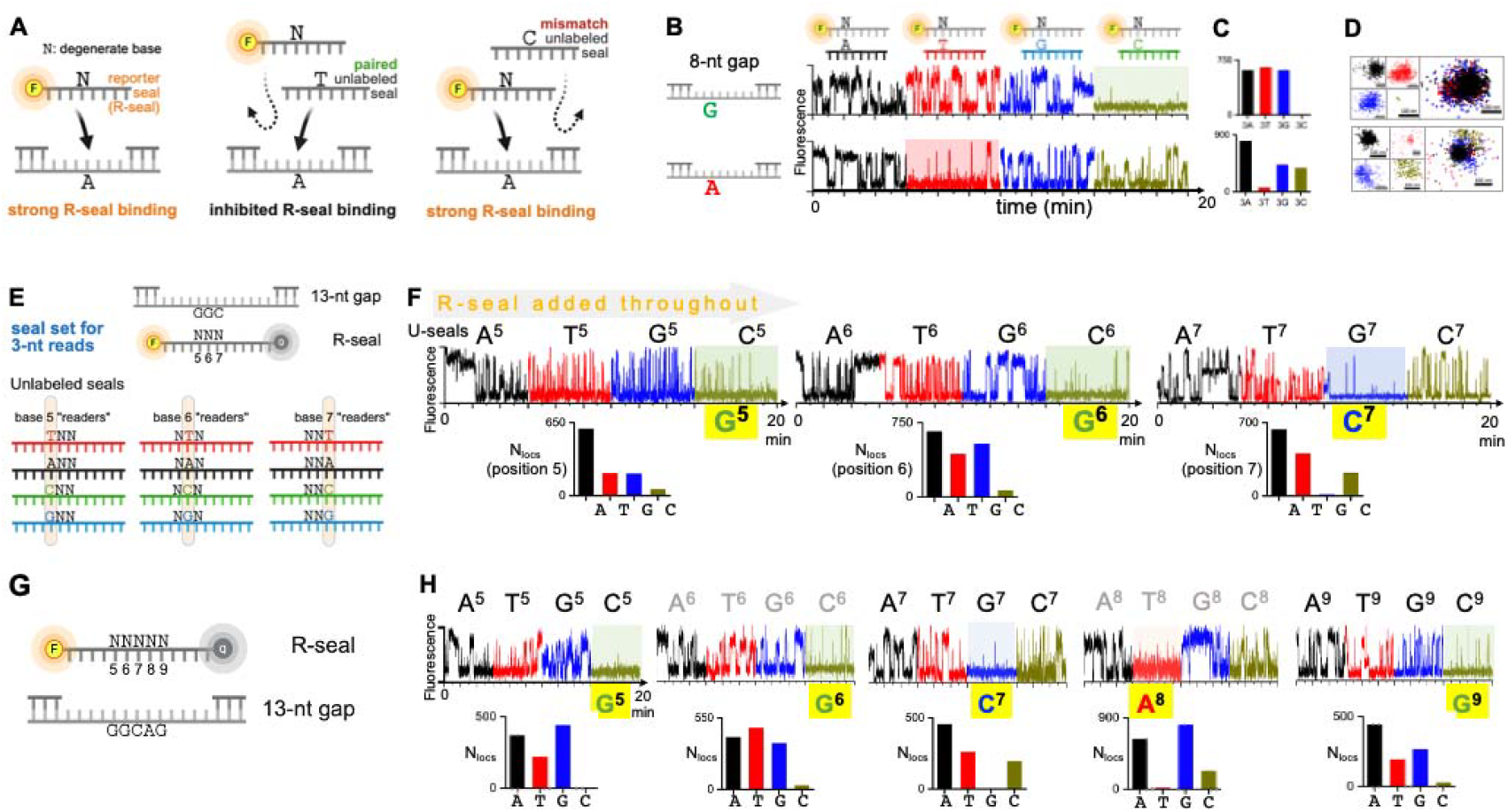
Single-molecule sequencing via competitive inhibition of transient hybridization. **(A)** Concept. Left: a labelled seal (R-seal; containing a degenerate base N) binds transiently to a gap and gives a fluorescence signal. Middle: in the presence of a complementary (“paired”) unlabeled seal, R-seal binding is reduced, and fluorescence is suppressed. Right: in the presence of a mismatched unlabeled seal, R-seal binding is maintained, and fluorescence is observed. **(B)** Reading single nucleotides. Top row: examining gap position 3 in Gap8-G^3^ using 200 nM of labelled R-seal and a set of 4 unlabeled U-seals (added sequentially, all at 600 nM); the complementary U-seal (U8-C^3^, in green) suppress R-seal binding, enabling base identification. Bottom row, same as above but for Gap8-A^3^. **(C)** Counts of seal localizations. The count for the complementary U-seals is very low compared to the three 1-nt mismatch seals. **(D)** Left: mapped localizations from single U-seals. Right: superimposed localizations from all 4 seals. **(E)** DNA design for reading a 3-base sequence (here, G^5^G^6^C^7^) using competitive inhibition. For details, see main text. **(F)** Demonstration of ability to read three bases using R-seal/U-seal combinations. R-seal (R13-N^5^N^6^N^7^) concentration was kept at 500 nM throughout all interrogations. Each of the three bases was evaluated on a different surface. A single base is read via a series of 4 sequential interrogations with 4 U-seals (1.5 μM); here, we refer to each seal using only the identity and position of the interrogating U-seal base (e.g., “A^5^” instead of “U13-A^5^N^6^N^7^”). Only complementary U-seals substantially decrease R-seal binding (highlighted traces). Bottom, number of localizations for each seal interrogation. **(G)** DNA design for sequencing a 5-base sequence (here, G^5^G^6^C^7^A^8^G^9^) using competitive inhibition. **(H)** Demonstration of ability to read five bases. Experiments as for 3-base sequencing apart from that the R-seal (R13-N^5^N^6^N^7^N^8^N^9^) and U-seals were both added at 1 and 2 μM, respectively. Results shown as in panel (F).

### Sequencing of up to 5 bases

Sequencing 3 bases using competitive inhibition required increasing both the number of degenerate bases in the R-seal (from 1 to 3) and the gap length (to 13 nt; **Fig. 3E**). Adding degenerate bases decreases the concentration of the fully complementary seal (e.g., for seals with 3 degenerate bases, only one in 64 sequences is fully complementary), effectively lowering the hybridization on-rate. To maintain frequent but short R-seal binding to the gap, and maximize the assay contrast, we adjusted our R-seal and U-seal concentrations, added a quencher to the R-seal to suppress fluorescence of unbound seals, and modified our hybridization buffer (**figs. S10-12** and *Methods*). We also modified our U-seals to include degenerate bases; e.g., for reading position 5 within a gap with unknown sequence at 5, 6 and 7 (Gap8-N^5^N^6^N^7^), we used 4 U-seals: U13-A^5^N^6^N^7^, U13-C^5^N^6^N^7^, U13-G^5^N^6^N^7^, and U13-T^5^N^6^N^7^ (**Fig. 3E**).

To read a base at position 5, we performed four sequential interrogations using mixtures of the R-seal and each of the four U-seals. Our results showed that R-seal binding decreases only with U13-C^5^N^6^N^7^ (abbreviated as “C^5^”; **Fig. 3F**, green trace), and that we can read a G at gap position 5 with ~96.4% accuracy (240 out of 249 molecules). We obtained similarly high accuracies for gap positions 6 and 7 (~96.5%; **Fig. 3F** and **fig. S13; table S2**), successfully identifying all three bases. To establish our ability to read multiple bases sequentially, we also sequentially interrogated a three-base sequence (GGC) on the same set of immobilized DNA gap molecules (**fig. S13B**), achieving ~99% base-calling accuracy (188 out of 189 molecules).

We also examined a 5-base sequence by using the same 13-nt gap, increasing the number of degenerate bases in the R-seal and U-seals (to 5 and 4, respectively; **Fig. 3G**), and adjusting the R-seal and U-seal concentrations (**figs. S14-15**). For gap position 5, only seal U13-C^5^N^6^N^7^N^8^N^9^ (abbreviated as “C^5^”) substantially decreased R-seal binding (**Fig. 3H** and **fig. S16**), detecting the G base with ~95% accuracy (195 out of 205 molecules). We achieved similar accuracies (~97%) for the remaining 4 positions (**Fig. 3H** and **fig. S16; table S2**).

Our results show that we can sequence up to 5 bases with high accuracy, paving the way for sequencing 4^5^=1,024 sequences on the same surface (requiring 4*5 = 20 interrogation rounds), and linking them to single-molecule function.

### Linking protein-DNA interactions to DNA sequence for single molecules

To link single-molecule phenotypes to sequence, we examined DNA interactions of transcription factor catabolite activator protein (CAP (*18*); **Fig. 4A**). In the presence of cyclic adenosine monophosphate (cAMP), CAP binds with high affinity to consensus sequence 5’-AAA<u>TGTGA</u>TCTAGA<u>TCACA</u>TTT-3’ (CAPcons; (*19*)), with the underlined sequences controlling specificity; **Fig. 4B**).

**Figure 4.**
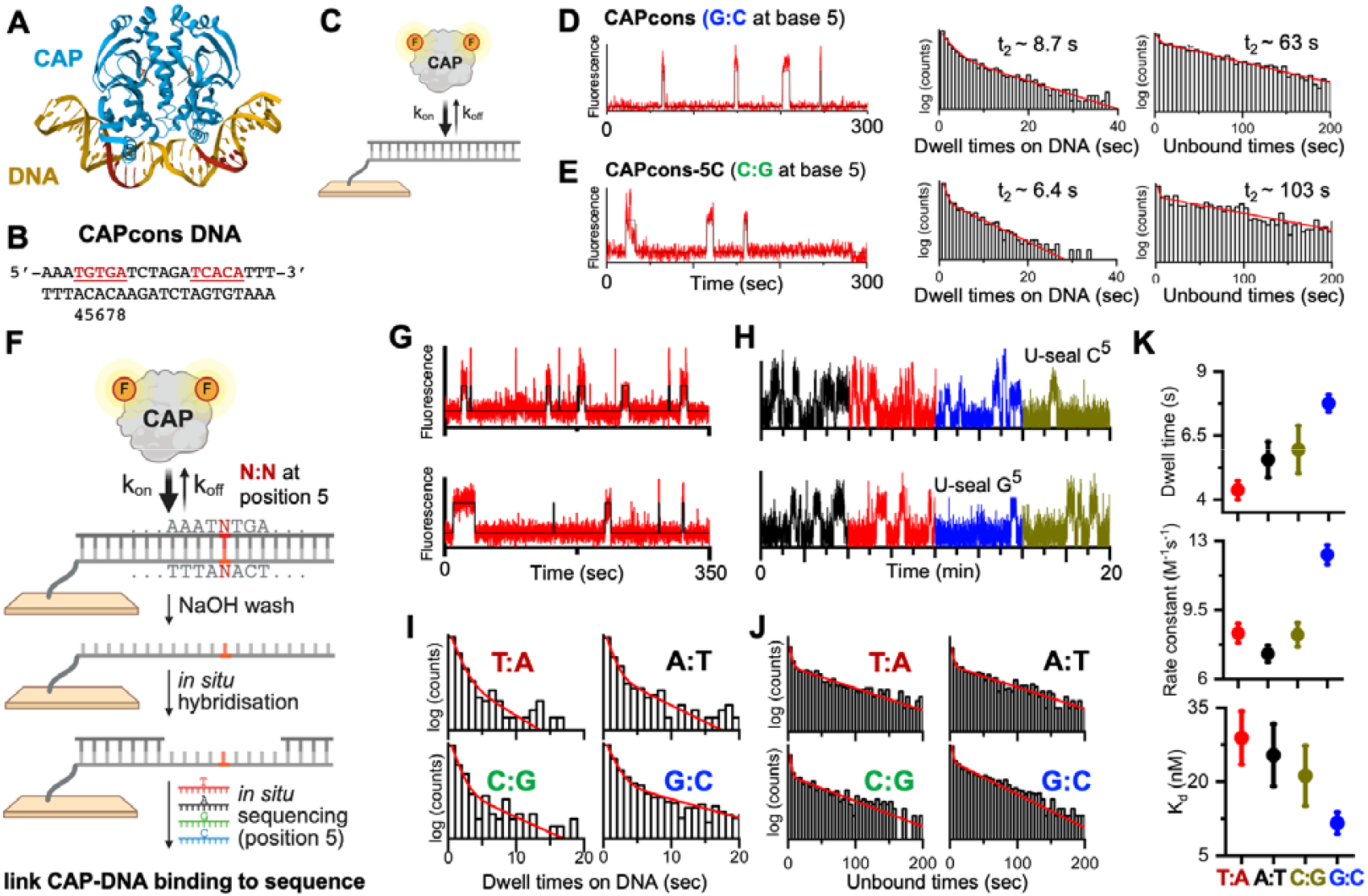
Linking kinetics of protein-DNA interactions to DNA sequence: transcription factor binding to its cognate DNA. **(A)** Structure of *Escherichia coli* catabolite activator protein (CAP) bound to its consensus DNA site (1CGP); red segments denote the 5-bp sequences conferring sequence specificity. **(B)** CAPcons sequence. (**C)** Labelled CAP binds to DNA variants transiently, yielding on- and off-rates. **(D)** Left: CAP binding to CAPcons DNA. HMM analysis identifies a high-intensity state (CAP-DNA complex), and a low-intensity state (free DNA). Right: dwell-time and off-time distributions fit best to double-exponential distributions; the longer lifetimes (t_2_) identify the lifetimes for stable binding and the frequency of stable complex formation. **(E)** As for (D), but for CAPcons variant with C:G at position 5. (**F)** Linking kinetic phenotype to sequence. After evaluating CAP binding to surface-immobilised CAPcons variants, the DNA is denatured, and gapped-DNA is formed via *in situ* hybridisation of two flanking strands; gap position 5 is then read. (**G)** CAP binding to variant CAPcons library. **(H)** Gap interrogation of the two DNA molecules in panel (G) using competitive inhibition; the variants are identified as 5G (top) and 5C (bottom). **(I)** Dwell-time histograms for bound CAP for CAPcons variants. (**J)** Dwell-time histograms for unbound CAP for CAPcons variants. **(K)** Dwell times, association rate constants k_on_, and K_d_ values for CAPcons variants; error bars: S.E.M. from 2 experiments.

To observe CAP-DNA binding, we surface-immobilized CAPcons DNA, added labelled CAP, and observed transient CAP binding to DNA (**Fig. 4C** and **Fig. 4D**, left). CAP dwell times on DNA (**Fig. 4D**, middle; **table S3**) showed a lifetime t_2_ ~8.7 s, which reflects stable CAP binding to CAPcons DNA; this assignment was supported by control experiments in the absence of DNA or cAMP (**fig. S17**). Binding-frequency analysis (using unbound-time distributions; **Fig. 4D**, right) yielded an association rate constant k_on_ ~1.3 × 10^7^ M^−1^s^−1^ and a K_d_ ~9.1 nM. A variant featuring a C:G pair at position 5 (vs. the consensus G:C) showed weakened CAP-DNA interactions (t_2_ ~6.4 s; k_on_ ~7.7 × 10^6^ M^−1^s^−1^; **Fig. 4E** and **table S3**), and 2-fold reduced affinity (K_d_ ~20 nM).

We then linked CAP-DNA binding for a *single* DNA molecule to the sequence of the *same* DNA molecule (**Fig. 4F**) for a library of surface-immobilized CAPcons DNA molecules, where position 5 of CAPcons was randomized. In step 1, we evaluated CAP-DNA binding as above. In step 2, the dsDNA was denatured, leaving only the biotinylated bottom DNA strand on the surface; subsequently, two DNA strands were hybridized efficiently *in situ* (**fig. S18**) to form a surface-immobilized gapped DNA with a 9-nt gap matching the first 9 bases of CAPcons (5’-AAA<u>TGTGA</u>T-3’).

In step 3, we analyzed the same field-of-view and sequenced position 5 using competitive inhibition. Using the surface locations of each protein-DNA binding and sequencing event, we linked CAP-DNA binding events (from step 1; **Fig. 4G**) to sequencing traces for single molecules (**Fig. 4H** and **fig. S19**). Binding analysis showed that G:C (known to form 2 hydrogen bonds from guanosine N7 and O^6^ atoms to CAP Arg180 side chain (*18, 20*)) leads to a slower off-rate and a faster on-rate relative to the other variants (**Fig. 4I-K** and **table S3**), leading to a K_d_ 1.9- to 2.6-fold lower than the non-consensus variants, corresponding to a ~0.6-0.9 kT (~0.35-0.55 Kcal/mole) stabilization of the complex versus non-consensus variants. Our results reproduced those from pure sequences (**Fig. 4D-E**) and established our ability to work with labelled proteins and link protein-DNA interactions with DNA sequence.

### Sequence-dependence of initial transcription

To test whether SPIN-Seq can address complex non-equilibrium reactions, we applied it to initial transcription, largely controlled by sequence-specific interactions between gene promoters and RNA polymerase (RNAP). Single-molecule studies (*21–23*) showed that, during bacterial transcription, RNAP pauses after synthesizing a 6-mer RNA, and enters a branched pathway (**Fig. 5A**) involving a productive path (where the 6-mer is extended to full-length transcripts), an abortive path (where short RNAs dissociate and synthesis restarts), or a “futile-cycling” path (where the 6-mer fluctuates between backtracked and non-backtracked states (*22*)). The initial transcribed sequence affects initiation pausing (*22*); next-generation sequencing (NGS) work on sequence libraries (*24*) identified a pause-inducing motif, TNYG (Y: pyrimidine), operational when the A-site of the RNAP active center is at positions +3,+4,+5 and +6 from the transcription start site (*25, 26*). However, the sequence-dependence of initiation-pausing kinetics and path branching during initial transcription were unknown.

**Figure 5.**
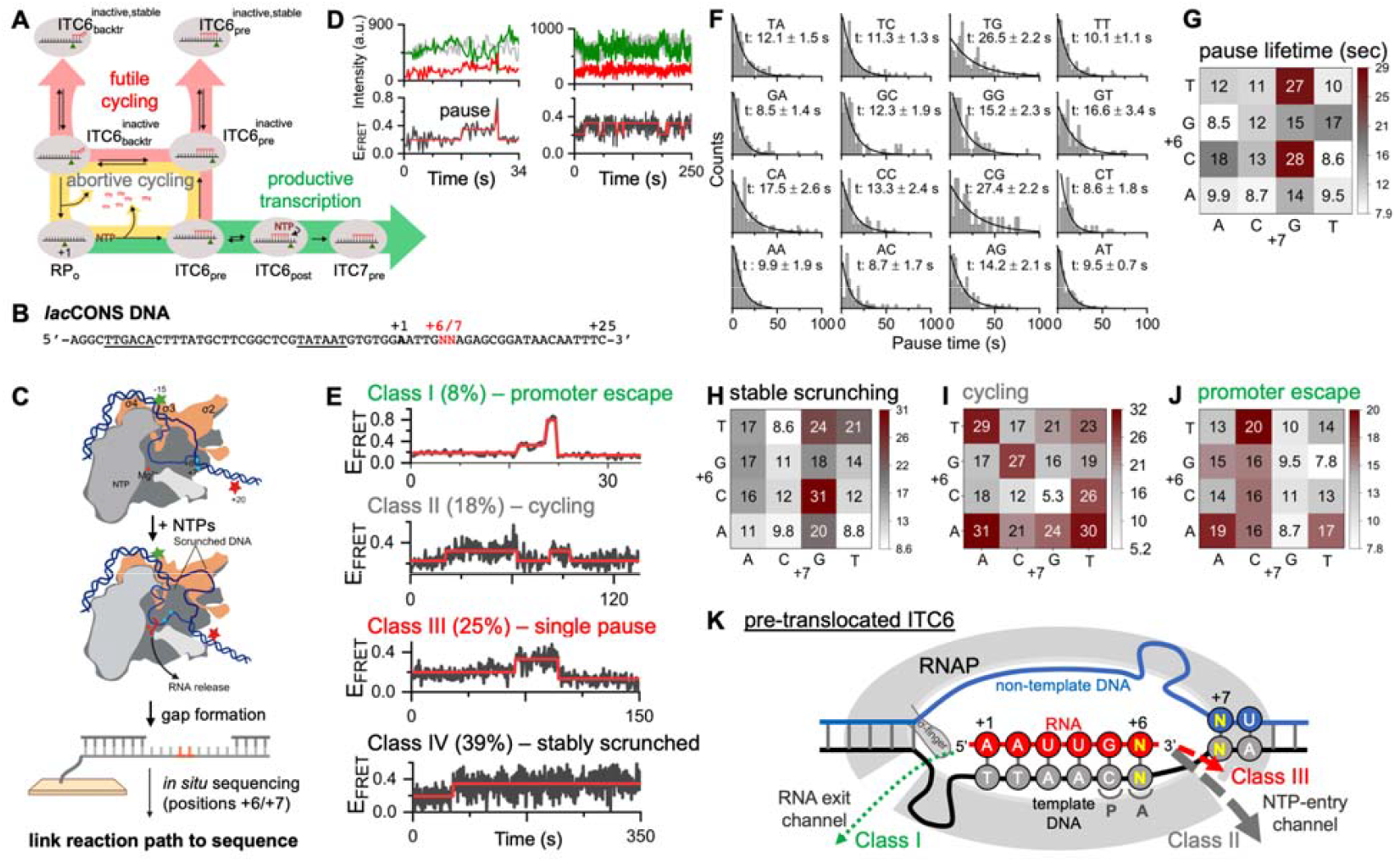
Linking kinetics of protein-driven reactions to DNA sequence: initial transcription mechanism. **(A)** Schematic of the 3 main paths for the short RNAs synthesized during initial transcription. Green path: productive RNA extension and promoter escape to elongation. Yellow path: abortive RNA release and restart of RNA synthesis. Pink path: futile cycling between a stably scrunched and RNA-backtracked state; RNAP tends to pause at ITC6 (initially transcribing complex with 6-nt RNA). **(B)** Sequence of the non-template strand of *lac*CONS promoter 16-variant library, generated by randomizing positions +6 and +7 (in red). Underlined sequences: −35 and −10 promoter elements. **(C)** Transcription complexes on the 16-variant library. After NTP addition to open complexes (top), transcription leads to FRET changes due to movement of the DNA downstream of the transcription start site (middle). After recording FRET changes, protein and template strand are removed, and positions +6 and +7 are read using competitive inhibition (bottom). **(D)** Examples of anticorrelated donor (green) and acceptor (red) fluorescence changes, consistent with single-molecule FRET fluctuations. Red line in FRET trace: HMM-based states and transitions. **(E)** Examples of traces belonging to the 4 main functional classes of active transcription complexes (*Main Text* for descriptions). **(F)** Distribution of initiation-pause lifetimes, and single-exponential decay fitting to obtain the pause lifetime for each variant. **(G)** Heatmap of pause lifetimes for different sequences. **(H-J)** Heatmap of propensity for stable-scrunching (H), cycling (I) and promoter escape (J) for different sequences, shown as fraction of molecules in the population belonging to this class. **(K)** Schematic of the pre-translocated ITC6 state. The 6-mer RNA (in red) can overcome the barrier of σ-finger to escape to elongation (Class I events), backtrack into the NTP-entry channel and dissociate (Class II events), or backtrack and remain arrested in the NTP-entry channel (Class III events). Class IV events involve an inactive state of ITC6, likely reflecting an inactive RNAP conformation. P and A: P-site and A-site of the RNAP active center.

To examine how DNA sequence affects initiation pausing, we generated a library of 16 *lac*CONS promoter variants (**Fig. 5B**) with degenerate bases at positions +6 and +7 (*Methods*; **fig. S20**). To monitor pausing, we applied an existing single-molecule FRET strategy (*21*) by labelling each variant with a donor at position −15 on the non-template strand, and an acceptor at position +20 on the template strand (**Fig. 5C**). As the acceptor approaches the donor during initial transcription (due to DNA scrunching (*21, 27*)), FRET reaches an apparent FRET efficiency E_FRET_ of ~0.4, where pausing occurs upon synthesis of a 6-mer RNA. Further synthesis past the pause, and promoter escape leads to a transient E_FRET_ of ~0.6 before FRET returns to a low value (*21*).

To capture initial transcription kinetics, we formed RNAP-DNA open complexes on the variants, immobilized and imaged them (*Methods*); we observed a mean E_FRET_ of ~0.2, as in our previous work (**fig. S21A**). After adding 500 µM ApA and 200 µM of all NTPs, most molecules show FRET changes, indicating transcriptional activity (**Fig. 5D** and **fig. S21B**); these changes disappeared upon adding rifampicin (an antibiotic that blocks extension beyond a 3-nt RNA), confirming they were specific to transcription (**fig. S21C**).

After monitoring transcription for ~6.6 min, we removed the protein and template strand, annealed DNAs to create an 8-nt gap centered around positions +6 and +7, and sequenced them using competitive inhibition (**fig. S22**). We observed that 84% (1370 out of 1633) of the transcriptionally active molecules exhibited a pause at E_FRET_~0.4. These pausing molecules displayed 4 main classes of behavior (**Fig. 5E**). Class I molecules (14%) showed stepwise E* changes up to ~0.6 and decrease to a low FRET state, consistent with promoter escape and completion of transcription on the template (*21*). Class II molecules (21%) displayed extensive cycling between the ~0.2 and ~0.4 FRET states, suggesting abortive cycling, during which a short RNA (4- to 6-mer) is released from initial transcribing complexes (ITCs), allowing RNAP to restart RNA synthesis. Class III molecules (49%) showed only a single pause at E_FRET_~0.4, after which they return to ~0.2 and stay there for the rest of the trace; these molecules are likely to be inactive ITCs with backtracked RNA (*22, 23*). Finally, Class IV molecules (16%) reached the ~0.4 FRET state and stayed there for the rest of the trace (~5 min); these molecules are stably scrunched complexes, major intermediates during *lac*CONS initial transcription (*21, 22*). The convolution of all signals is visualized in heat maps of FRET temporal evolution as a function of sequence, providing an overview of differences between sequences (**fig. S23**).

To determine the pause lifetimes of all 16 variants, we measured the duration of each dwell at E_FRET_~0.4 for Classes I-III (**Fig. 5E**), plotted the first-pause dwell-time distributions for each sequence and determined pause lifetimes (**Fig. 5F**). Consistent with earlier work (*22, 24*), sequence T^+6^G^+7^, found both on natural *lac* promoter and synthetic *lac*CONS promoters, showed a long pause (~27 s, n=110). Sequence C^+6^G^+7^ led to similar pausing (~27 s, n=80), while G^+6^A^+7^ showed the shortest pause (~8.5 s, n=88); hence, just a 2-nt sequence can lengthen the pause by >3-fold. In general, promoters with pyrimidines (Y; C or T, the preference being stronger for C) at +6, and a G at +7 exhibited long pausing, supporting an initiation pause element of (C/T)^+6^G^+7^ or Y^+6^G^+7^ (**Fig. 5G**). This sequence matches the consensus for pausing in transcription elongation (*25, 26*), pausing in initial transcription (*24*), and σ-dependent pausing in early elongation (*28*), as determined using NGS; in all cases, the pause-inducing sequence preferences reflected the tendency of specific nucleotides to disfavor RNAP translocation from its pre- to post-translocated state, with Y^+6^ preferences reflecting RNAP active-site binding preferences for specific nucleotides, and G^+7^ preferences reflecting effects of duplex stability and active-site binding preferences (*26, 29*).

To verify our findings, we prepared the C^+6^G^+7^ and A^+6^T^+7^ variants, associated with long and short pausing, respectively. Analyzing the variants separately recapitulated the findings from the library, validating our approach (**fig. S24**).

We also examined how sequence impacts the transcription profile by looking at the sequence dependence of the tendency to belong to a given class (**Fig. 5H-J** and **fig. S23**). Sequences T^+6^G^+7^ and C^+6^G^+7^ (showing the longest pauses in Classes I-III) were linked to stable scrunching (Class IV; **Fig. 5H**), strongly suggesting that Y^+6^G^+7^ sequences favor formation of a common reaction intermediate that precedes entry into either a long but transient paused state, or an exceptionally long paused state. The latter intermediate is likely an inactive pre-translocated ITC6 (**Fig. 5A**), the branchpoint entry to Class II, III, and IV molecules.

In contrast, cycling molecules (Class II) show a striking preference for A/T at positions +6 and +7 (e.g., 31% of A^+6^A^+7^ traces cycle, where only 5% of C^+6^G^+7^ traces cycle; **Fig. 5I**). It has been proposed (*21, 22*) that cycling molecules leave the paused state via RNA backtracking, RNA entry in the NTP-entry channel, RNA release, and restart of RNA synthesis (**Fig. 5K**). Our results strongly support this proposal, since the observed A/T preference indicates low duplex stability and easier opening of the RNA-DNA hybrid at the transcript 3’-end, matching a well-known determinant for RNA backtracking (*30, 31*). Notably, many complexes with A/T sequences at positions +6 and +7 will synthesize a 7-mer (since they pause for shorter time), and have a 2 A/U nt at the RNA 3’-end, while still encountering the σ^70^ finger (a structure blocking the path of growing RNA, shown not to be fully displaced on lacCONS until 10-nt RNAs are synthesized (*32*)). Efficient cycling also requires that short RNAs dissociate rapidly from the NTP-entry channel to allow synthesis to restart; consistent with this, A/T sequences at positions +6 and +7 are under-represented in single-pause events (Class III; **Fig. 5J** and **fig. S25**), which reflect RNA backtracking into the NTP-entry channel and transcriptional arrest (**Fig. 5K**).

Finally, promoter-escape events (Class I) disfavor the presence of G at +7, while showing a slight preference for a C at +7, with T^+6^C^+7^ leading to the most productive transcription (**Fig. 5J**). This preference reflects the fact that “productive” sequences must avoid the long pausing and stable scrunching favored by a G at +7, and the abortive cycling favored by A/T at +6/+7; as such, our analysis can offer guidance for promoter design.

### Conclusions

We introduced SPIN-Seq, a method that leverages protein-free *in situ* sequencing to link functional and structural phenotypes of single DNA molecules with their sequence. The sequencing aspect of the method relies on direct and highly accurate detection of single DNA mismatches via transient hybridization. As we show by analyzing the interaction of a transcription factor with its cognate DNA site, SPIN-Seq uses standard surfaces for single-molecule imaging of labeled proteins, making it applicable to a wide repertoire of DNA-interacting biomolecules, including proteins, nucleic acids, and small molecules. The method quantifies DNA interactions across a broad range of affinities (sub-nM to 10 μM), operates without amplification or commercial sequencing, and is compatible with diverse libraries of nucleic acids and DNA-encoded molecules, such as aptamers and DNA-encoded drugs.

Our work demonstrates that even a modest library of 16 DNAs offers rich mechanistic insight into a complex non-equilibrium reaction, showing directly that just two nucleotides can dramatically alter transcription initiation pausing and the distribution between productive and non-productive paths. We also reveal that initial transcription pausing has further similarities with elongation pausing, with additional tuning contributed by initiation factors and by different extents of DNA melting and scrunching. Given the homology of bacterial and eukaryotic RNAPs, sequence-dependent branching may also shape eukaryotic transcription, where a finger-like element in TFIIB causes initiation pausing at positions +7 to +9 (*33*), complementing regulation through promoter-proximal pausing (*34*).

The power of SPIN-Seq will increase further by extending it to 5-nt sequencing, enabling DNA libraries of over a thousand members, and by boosting assay throughput with automated microfluidics (*10, 11*), larger fields of view (*35*), and higher molecular densities enabled by high-precision localization ((*13, 36*); **Fig. 3D**). SPIN-Seq can also be integrated with other single-molecule approaches that incorporate fluorescence, broadening opportunities for biophysical analysis. Together with recent single-molecule multiplexed methods (*10, 11, 37–39*), SPIN-Seq offers a powerful framework to systematically decode how DNA sequence shapes the dynamics of molecular interactions.

## Acknowledgements

We thank C. Hepp for help with the use of fiducial markers for drift correction, S Chatzimichail for help with microfluidics, A. Shivalingam for help with DNA synthesis, and the MICRON Advanced Bioimaging Facility (supported by Wellcome Strategic Awards 091911/B/10/Z and 107457/Z/15/Z) for access to a Nanoimager microscope in their facilities.

## Author contributions

Conceptualization: A.N.K. Methodology: J.P.H., R.A., P.T., Q.Z., H.S., M.K., A.N.K. Investigation: J.P.H., R.A., P.T., A.H.E.S., A.M., H.E.S, A.N.K. Software: P.T., Q.Z., H.S. Formal analysis: J.P.H., R.A., P.T., Q.Z., A.N.K. Visualization: J.P.H., R.A., Q.Z., A.N.K. Funding acquisition: J.P.H., R.A., T.B., A.N.K. Project administration: A.N.K. Supervision: T.B., A.N.K. Writing – original draft: J.P.H., R.A., A.N.K. Writing – review & editing: J.P.H., R.A., A.N.K.

## Data and materials availability

Single-molecule movies and fluorescence intensity time-traces will be available upon request; large datasets will also be deposited to a long-term public repository before publication. Software for image analysis and base calling are available on Github; see SI for links. Correspondence and request for materials and datasets should be addressed to A.N.K.

## Supplementary Materials

Materials and Methods

Supplementary Text

Figs. S1 to S25

Tables S1 to S3

References (*40–49*)

