## Supplemental table and figures for "Unraveling single-molecule reactions via multiplexed in-situ DNA sequencing"

**The PDF file includes:**

Materials and Methods

Supplementary Text

Figs. S1 to S25

Tables S1 to S3

References

Materials and Methods

Preparation of DNA fragments**.** A complete list of all DNA fragments is found in **table S1**. Oligodeoxyribonucleotides (“oligos”) used in **Fig. 2** were synthesized on an Applied Biosystems 394 automated DNA/RNA synthesizer using a standard 1.0 μmole phosphoramidite cycle of acid-catalyzed detritylation, coupling, capping, and iodine oxidation. Stepwise coupling efficiencies and overall yields were determined by the automated trityl cation conductivity monitoring facility and was >98.0%. All β-cyanoethyl phosphoramidite monomers were dissolved in anhydrous acetonitrile to a concentration of 0.1 M immediately prior to use with coupling time of 50 s for normal A, G, C, and T monomers and was extended to 600 s for 5’-amino, 5’-biotin and 5-amino C6 dT phosphoramidite monomers (from Link Technologies Ltd.). 5’-Amino and internal 5-amino C6 dT modified oligonucleotides on resin were treated with a solution of diethylamine (10% in acetonitrile) for 20 min to selectively remove the cyanoethyl protecting groups. Cleavage and deprotection of oligonucleotides were achieved by exposure to concentrated aqueous ammonia solution for 60 min at room temperature followed by heating in a sealed tube for 5 hrs at 55°C.

Oligonucleotides labelled with either an ATTO647N or Cy3B were prepared by NHS-ester coupling. The unlabelled pre-cursor oligonucleotide (40 nmol) was freeze-dried and resuspended in sodium bicarbonate buffer (18 μl, 0.5 M, pH 8.5). Separately, Cy3B or ATTO647N NHS-ester (6 μl, 20 nmol/μl in DMSO) was mixed with DMSO (6 μL). The NHS-ester solution was then added to the oligonucleotide solution and mixed vigorously before shaking (800 rpm, room temperature). After 3 hrs, reactions were diluted with water (970 μl) and purified by desalting on a disposable gel-filtration column (NAP10, GE Healthcare Life Sciences, cat. no. GE17-0854-02) according to the manufacturer’s instruction to remove most of the free dye. Labelled oligonucleotides were further purified by reverse-phase HPLC using a Gilson HPLC system with ACE® C8 column (10 mm x 250 mm, pore size 100 Å, particle size 10 μm) with a gradient of buffer A (0.1 M TEAB, pH 7.5, where TEAB = triethylammonium bicarbonate) to buffer B (0.1 M TEAB, pH 7.5 containing 50% v/v acetonitrile) and flow rate of 4 ml/min over a total run time of 17.5 min. For all ATTO647N-labelled oligonucleotides, a gradient of 70–100% buffer B was used. For Cy3B-labelled oligonucleotides, a gradient of 20–35% buffer B was used. The TEAB buffer and acetonitrile in the collected fractions were removed in vacuo or by freeze drying.

All oligonucleotides were characterised by negative-mode electrospray using a UPLC-MS Waters XEVO G2-QTOF mass spectrometer and an Acquity UPLC system with a BEH C18 1.7 μm column (Waters).

Oligonucleotides used in all remaining figures were procured from Metabion and Biomer corporation. Commercially procured oligonucleotides were dissolved in nuclease-free water to a final concentration of 100 μM and stored at -20°C.

With regards to our DNA nomenclature (**table S1**), in a strand named Gap-(1-27)-Top^Cy3B,18^, “Gap” refers to the aim of the experiment (here, sequencing experiments with a gapped DNA); the term “(1-27)” refers to start and the end of the strand with respect to the 5' end, “Top” denotes the top strand, and the superscript (^Cy3B, 18^) specifies a Cy3B fluorophore attached to position 18. For easy of reference and brevity, we also use a shorter abbreviation for DNAs, which denotes the type of seal (or gap); the length of the seal (or gap); and the gap position and base interrogated. For example, “U8-A^3^” refers to a U-seal that is 8-nt long and has an A base at the position that corresponds to gap position 3 in the gap (i.e., interrogating for the presence of a T in the gap).

To form gapped-DNA substrates, as well as fully duplexed DNA substrates, oligonucleotides were annealed by mixing complementary strands in a ratio of 1:1 in hybridization buffer (200 mM Tris–HCl pH 8.0, 500 mM NaCl, 1 mM EDTA). The mixture was heated to 95°C for 5 min to denature the strands, followed by gradual cooling to 25°C at a rate of 2°C per min to facilitate annealing. The annealed DNA strands were then placed on ice until further use.

Preparation of CAPcons DNA library. To prepare the variant library for position 5 of the 22-bp CAPconsensus sequence (CAPcons), 1 µM of biotinylated 49-nt ssDNA having a degenerate base at position 25 (CAPcons-(1-49)-N^25^-B^Bio,49^) was annealed with 2 µM of complementary Cy3B-labelled 20-nt ssDNA (CAPcons-(1-20)-Top^Cy3B,11^). The top strand was then extended using Q5 DNA Polymerase (Thermofisher Scientific) and dNTPs (NEB) using melting, annealing and extension temperatures of 95˚C, 62˚C and 72˚C, respectively. The variant library was purified using QiAquick purification kit (QIAGEN). Success of the primer extension was verified by running the purified product in a 20% bi-acrylamide/acrylamide gel and imaged using a Typhoon gel scanning system. The band corresponding to the 49-bp dsDNA was excised from the gel and soaked in TE buffer overnight at room temperature. To concentrate the product, the DNA was precipitated by mixing the solution with 3X volume of ice-cold ethanol, 1 µL of RNA-grade glycogen (Thermo Fisher Scientific) and 20 mM sodium acetate. The mixture was kept at -20˚C for 4 hrs, followed by centrifugation and removal of the supernatant. The precipitated product was dissolved in 50 µL of assay buffer (AB: 20 mM HEPES pH 7.7, 80 mM MgCl_2_, 200 mM NaCl) and was stored at -20˚C. The final DNA concentration was measured using UV-Vis spectrophotometry on a Nanodrop ND1000 spectrophotometer.

Preparation of lacCONS DNA library. To prepare the variant library for positions +6 and +7 of *lac*CONS promoter (**fig. S20**), 2 µM of biotinylated and labelled fragment of the transcription non-template strand (lacCONS(-39/-10)-Top^Cy3B,-15; Bio,-39^) were first annealed with 1 µM of a fragment of the template strand having degenerate positions at positions +6 and +7 (lacCONS(-39/+10)-N^+6^N^+7^-B). The top strand was then extended using Q5 DNA polymerase (Thermo Fisher Scientific) and dNTPs (NEB), using melting, annealing and extension temperatures of 95˚C, 62˚C and 72˚C, respectively. A second fragment of the template strand (lacCONS(+11/+49)-B^Atto647N,+20^) was then annealed to a second fragment of non-template strand (lacCONS(+11/+49)-Top), and the annealed strand was ligated to the primer-extended product using 0.5 µL of 2,000,000 units/ml T4 DNA ligase (NEB) and incubating overnight at 16˚C. Success of the primer extension was verified by running the purified product in a 20% bi-acrylamide/acrylamide gel and imaged using a Typhoon gel scanning system. The band corresponding to 86-bp sequences containing both Cy3B and ATTO647N fluorophore was excised from the gel and soaked in TE buffer for overnight. The gel-extracted product was precipitated by mixing the solution with 3X volume of ice-cold ethanol, 1 µL of glycogen, and 20 mM sodium acetate, and keeping at -20˚ C for 4 hrs. The precipitated DNA was finally dissolved in 50 µL of AB, and stored at -20˚C.

Preparation of labelled CAP derivative (Alexa647-CAP). We used a CAP derivative with two reactive Cys residues (Cys17; Ser178), that has been extensively characterized in terms of function and labelling (*40*, *41*). CAP overexpression and purification was performed using plasmid pHSCRP-His6-H17CC178S [constructed from plasmid pAKCRP-His6 (*42*) using site-directed mutagenesis (Q5 Site-Directed Mutagenesis Kit, NEB)]. Fluorescence labelling was performed by incubating 20 μM CAP-(Cys17; Ser178) with 400 μM Alexa647 maleimide in buffer A [40 mM HEPES-NaOH pH 7.3, 100 mM NaCl, 2 mM TCEP and 5% glycerol] 12 hrs on ice. The fluorescent probe labelled protein (Alexa647-CAP) was purified using a G25 Sephadex gel filtration column pre-equilibrated with buffer B (40 mM Tris-HCl pH 8, 100 mM NaCl, 0.1 mM TCEP and 5% glycerol), followed by a purification step on Ni:NTA agarose affinity column (and elution with 1M imidazole in buffer B), followed by buffer exchange (5 cycles of concentration to 0.5 ml volume and dilution with 5 ml buffer B) using a 3 kDa MWCO Amicon Ultra-15 centrifugal ultrafilter (Millipore). The purified Alexa647-CAP was stored in aliquots at -80^o^C. The labelling efficiency was 40%.

Preparation of RNA polymerase holoenzyme. *Escherichia coli* RNAP holoenzyme was expressed and purified by co-expressing genes of RNAP β, β’, α, ω and σ^70^. A primary culture of 20 mL LB broth was inoculated using a single colony of BL21 DE3 *E. coli* strain transformed with plasmid pV10 (*43*) and plasmid pRSFduet-sigma (*44*). Ampicillin (100 μg/ml) and kanamycin (50 μg/ml) were used as selection antibiotics. A second culture (2*1L) was started using 10 mL of primary culture using the same antibiotic level and was grown at 37˚C until O.D. reached 0.6. Inducer IPTG was added at 1 mM and the cultures were further incubated at 37˚C for 3 hrs. Cells were harvested and lysed followed by precipitation of cell lysate using polyethyleneimine and ammonium sulphate. Purification of the protein was performed using Ni-NTA agarose affinity column with elution using 200 mM imidazole in buffer A. Affinity purified sample was passed through anion-exchange chromatography equipped with Mono-Q 10/100 GL. Fractions containing the protein of interest were pooled together and concentrated using a 30 kDa MWCO Amicon Ultra-15 centrifugal ultrafilters.

Preparation of surface-immobilised DNA for imaging. Microscope coverslips and slides were prepared essentially as described (*45*). Briefly, borosilicate coverslips (#1.5 Menzel/ThermoFisher) were first burned in a furnace at 500^o^C, and were plasma-cleaned; subsequently, the coverslips were treated sequentially with acetone, 2% Vectabond aminosilane (Vector Labs) in acetone, and water, dried under nitrogen, and bonded to 6-mm silicone gaskets (GBL103280; Grace Bio-Labs), creating observation wells with aminosilane-functionalized glass. Wells were coated with 20 µl of 30 mM mPEG-SVA (Laysan Bio) and 0.75 mM Biotin-PEG-SVA (Laysan Bio) in MOPS-NaOH (pH 7.5) for 90 min at 22°C, washed with PBS, then treated with 30 μl of 10 μM NeutrAvidin (ThermoFisher) in 0.5xPBS for 10 min at 22°C. After PBS washes, wells had NeutrAvidin-biotin-PEG/mPEG-functionalized glass floors. The biotinylated DNA constructs were added to the observation wells to a final concentration of 50 pM, and were incubated until achieving an immobilisation density of ~0.2 molecules per mm^2^, and then were washed with AB. Next, we added to the observation wells 40 µL of FluoSpheres™ (Biotin-Labeled Microspheres, 0.2 μm, yellow-green fluorescent (505/515), 1% solids) diluted to 1:2000 volume in AB as fiduciary markers, and incubated for 20-40 s. We then washed the wells three times with AB, added 20 µL of 50 pM biotinylated gapped-DNA constructs in imaging buffer 1 (IB1: 20 mM HEPES pH 7.7, 200 mM NaCl, 80 mM MgCl_2_, 6 mM BSA, 1 mM TROLOX, 1% Glucose, 40 µg/ml catalase and 0.1 mg/ml glucose oxidase), incubated for 10-30 s, until achieving an immobilisation density of 0.2 molecules per µm^2^ on an average and washes three times with 200 µL of AB.

To surface-immobilise DNAs for single-base sequencing in a mixed population with prior information (**Fig.** **2E**), four different 8-nt DNA-gapped molecules were sequentially immobilized on the surface at a concentration of 50 pM. First, we immobilized Gap8-A^3^ (prepared by annealing Gap-(1-65)-A^30^-B^Bio,1^, Gap-(1-27)-Top^Cy3B,18^, and Gap-(36-65)-Top) and registered the locations of the DNA molecules by exciting with 0.2 mW of 532 nm laser, followed by photobleaching using 532 nm laser at 2 mW power. We then immobilized Gap8-T^3^ (prepared as above but using Gap-(1-65)-T^30^-B^Bio,1^ as a bottom strand) registered the DNA locations, and photobleached the fluorophores. We repeated the same procedures for Gap8-G^3^ and Gap8-C^3^. We then interrogated the gapped molecules using 4 rounds of transient seal hybridization.

Single-molecule imaging. All single-molecule measurements were performed using a Nanoimager-S microscope (Oxford Nanoimaging), equipped with 100x 1.4 NA objective, and a sCMOS camera (Orca Flash4 v3). The emission channel was split into two channels, one corresponding to wavelengths 498-551 nm and 576-620 nm, and the other corresponding to a 685/40 nm band (using 640LP channel splitter dichroic). An objective-based total internal reflection fluorescence (TIRF) illumination mode at an angle of 56˚ was used. All sequencing-related imaging experiments were carried out with continuous-wave excitation of 532 nm laser at 0.2 mW power, and 638 nm laser at 0.6 mW power, using a field-of-view of approximately 80 μm x 50 μm. For the experiments in **Fig. 2A-B**, 100-ms exposures were used; for experiments in Fig. **2C-2D**, the localization image was collected using 200-ms exposures, and the sequence-interrogation movies using 300-ms exposures for 1000 frames. For **Figs. 3-4** and **S5-19**, all movies (1200 frames) and localization images were collected using 250-ms exposures. For **Fig. 5**, and Figs. **S21-25**, 532 nm/638 nm alternating laser excitation and 200-ms exposures for 1000 frames were used.

Real-time in-situ sequencing assay. To call individual bases, we immobilized gapped-DNA molecules on the surface and registered their locations by exciting them with a 532-nm laser and detecting them using 200-ms exposures. In non-competitive approach of sequencing, 40 µL of 100 nM labelled seal was added on the surface in imaging buffer 2 (IB2: 20 mM HEPES, pH 7.7, 200 mM NaCl, 80 mM MgCl_2_, 5% dextran sulphate, 10% formamide, 1 mM TROLOX, 1% Glucose, 40 µg/ml catalase and 0.1 mg/ml glucose oxidase). The molecules were then excited using 532/638 nm laser excitation, and 1000 frames of 300-ms exposures were collected. Next, the surface was washed using 40 µL of AB for three times and incubated for 1 min with 40 µL of 100 nM of the labeled seal featuring a different standard base at the interrogated position; we then collected 1000 frames as described above. We repeated the process for the remaining two labeled seals.

Optimisation of sequencing by competitive inhibition. To optimize the seal concentrations for reading one base using competitive inhibition (**fig. S6**), we used an R-seal that comprises an 8-nt ATTO647N-labelled oligo with a degenerate base complementary to gap position 3 (R8-N^3^; **table S1**). Binding of the R-seal to an 8-nt gapped DNA (Gap8-G^3^, which features a G base at gap position 3, and was studied in **Fig. 2A**) leads to the appearance of fluorescent signals (**fig. S6**; see points where no U-seal was added).

To optimize the concentration of the U-seal complementary to the 8-nt gap (seal U8-C^3^, fully complementary to Gap8-G^3^), we monitored the gap-binding of 100 nM R8-N^3^ in the presence of 100-600 nM U8-C^3^ in imaging buffer 3 (IB3: 20 mM HEPES pH 7.7, 200 mM NaCl, 80 mM MgCl_2_, 10% Dextran Sulphate, 10% formamide, 1 mM TROLOX, 1% Glucose, 40 µg/ml catalase, 0.1 mg/ml glucose oxidase). Competition due to increasing U8-C^3^ leads to gradual loss of R-seal binding, as seen in fluorescence time-traces (**fig. S6A**), and in the markedly decreased R-seal dwell-times in the gap (**fig. S6B-C**), and in the markedly decreased mean number of R-seal localizations in the gap (**fig. S6D**). To optimize the concentration of the U-seal with 1-nt mismatch to the 8-nt gap (seal U8-T^3^), we monitored the gap-binding of 100 nM R-seal in the presence of 100-600 nM U8-T^3^ in IB3. R-seal binding also decreases with increasing U8-C^3^ (**fig. S7**) but to a much lower degree. We chose to use 100-200 nM R-seal and 300-600 nM U-seal (i.e., a 3:1 concentration ratio) for the competitive-inhibition assays, since they offer optimal contrast between complementary and 1-nt mismatched U-seals.

To optimize the seal concentrations for reading three bases using the competitive inhibition assay, we used a 13-nt gapped DNA (Gap13-G^5^G^6^C^7^, which features G, G, and C at gap positions 5, 6, and 7), prepared by annealing Gap-(1-65)-B^Bio,1^ with Gap-(1-27)-Top^Cy3B, 25^ and Gap-(41-65)-Top. We also used an R-seal that comprises a 13-nt ATTO647N-labelled oligo with three degenerate base complementary to gap positions 3, 6, and 7 (R13-N^5^N^6^N^7^; **table S1**). Since the 13-nt R-seal contains 3 degenerate bases (leading to the effective concentration of the exact complementary seal sequence in the mixture to be 64-fold lower than the total concentration used), we increased the R-seal concentration to 500 nM; this was possible using dark quencher BHQ1, which suppresses ATTO647N fluorescence until the R-seal binds to the gap (*46*). Since binding of a 13 nt R-seal will lead to longer dwells (which decrease sampling), we decreased the R-seal binding stability by increasing the formamide concentration. Upon testing R-seal binding in the presence of 10-25% formamide in IΒ3 (**fig. S10**), we identified 15% formamide as the optimal concentration for R-seal sampling.

To optimize the concentration of the U-seal complementary to interrogated position 5 in the 13-nt gap (seal U13-C^5^N^6^N^7^), we monitored the gap-binding of 500 nM R-seal in the presence of 0.5-3.5 μM U-seal in IB3 with 15% formamide (**fig. S11A**) and observed substantial decrease in R-seal binding at 0.5 μΜ U-seal and a more gradual decrease thereafter (**fig. S11B-D**). After performing an analogous experiment with the 1-nt mismatched U-seal (U13-T^5^N^6^N^7^; **fig. S12**), we observed that significant R-seal binding remains at 1.5 μM U-seal concentration. We thus chose to use 500 nM R-seal and 1.5 μM U-seal for the competitive inhibition assay, since they offer optimal contrast in R-seal binding between complementary and 1-nt mismatched U-seals.

To optimize the seal concentrations for reading five bases using competitive inhibition, we used the same gapped DNA as for 3-base sequencing. We also used an R-seal that comprises a 13-nt ATTO647N-labelled oligo with five degenerate base complementary to gap positions 5, 6, 7, 8 and 9 (R13-N^5^N^6^N^7^N^8^N^9^, **table S1**). Since the 13-nt R-seal contains 5 degenerate bases (leading to the effective concentration of the exact complementary seal sequence in the mixture to be 1024-fold lower than the total concentration used), we increased the R-seal concentration to 1 μM while maintaining the same buffer conditions. To optimize the concentration of the U-seal complementary to interrogated position 5 in the 13-nt gap (seal U13-C^5^N^6^N^7^N^8^N^9^), we monitored the gap-binding of 1 μΜ R-seal in the presence of 1-5 μM U-seal (**fig. S14A**) and observed substantial decrease in R-seal binding at 1 μΜ U-seal and a more gradual decrease thereafter (**fig. S14B-D**). After performing analogous experiments with the 1-nt mismatched U-seal (U13-T^5^N^6^N^7^N^8^N^9^; **fig. S15**), we observed that optimum contrast is achieved at 2 μM mismatch U-seal concentration. We thus chose to use 1 μM R-seal and 2 μM U-seal (a 1:2 concentration ratio) for the competitive inhibition assay.

Sample preparation for real-time CAP-DNA interactions. CAPcons DNA was prepared by annealing a 5’-biotinylated ssDNA (CAPcons-(1-49)-C^25^-B^Bio,49^) with its complementary ssDNA (CAPcons-(1-49)- G^25^-Top^Cy3B,13^) containing Cy3B at position 13; the Cy3B fluorophore allowed detection and localization of the DNA molecules on the surface. To prepare the G:C variant at position 5 of the CAPcons sequence, we altered the base at 25^th^ position of the biotinylated ssDNA sequence to G (CAPcons-(1-49)-G^25^-B^Bio,49^) and annealed with its complementary ssDNA sequence (CAPcons-(1-49)-C^25^-Top^Cy3B,13^).

To study CAP-DNA interactions in real-time (**Fig. 4B-C**), we immobilized one or more CAPcons variants on the surface by incubating 50 pM of 40 µL DNA on a PEGylated surface and washing three times with AB, followed by two washes with CAP-binding buffer (40 mM Tris pH 8, 100 mM KCl, 10 mM MgCl_2_, 5% glycerol, and 0.2 mM cAMP). We then added 1.25 nM AF647-CAP, and imaged binding in CAP binding buffer supplemented with 1 mM TROLOX, 1% Glucose, 40 µg ml^-1^ catalase and 0.1 mg ml^-1^ glucose oxidase. We used 250-ms exposures and collected 1800 frames.

In-situ DNA gap preparation and sequencing. To sequence the CAPcons variant library after the CAP-binding assay, we removed the non-biotinylated top strand, and created a gap around the positions of interest. We washed the surface three times using MQ water, incubated the surface with 20 mM NaOH for 30 s to de-hybridize the dsDNA, and washed three times washing with AB. To create a 9-nt gap around position 25, flanking strands CAPcons-(30-49)-Top and CAPcons-(1-20)-Top^Cy3B,11^ were incubated on the surface for 15 min at a final concentration of 2 µM in AB supplemented with 10% dextran sulphate.

To measure the efficiency of *in-situ* formation of DNA gaps, we first deposited the 8-nt gapped DNA used in **Fig. 2A** and localised the gapped DNA molecules (**fig. S18A**). After performing transient hybridization with the complementary seal, we removed the strands flanking the gap by washing the surface three times with MQ water, incubating it with 20 mM NaOH for 20 s, and washing it three times using AB. We reformed the DNA gaps *in situ* by incubating the surface with 100 nM of flanking strands Gap-(1-27)-Top^Cy3B,18^ and Gap-(36-65)-Top) in AB supplemented with 10% dextran sulphate for 15 min, and imaged the surface again (**fig. S18B**). Colocalization analysis between the initial localization image (**fig. S18C**) and the re-annealed image shows a ~70% colocalization (at a distance threshold of 7 pixels) of the initially deposited DNA gaps and the *in-situ* formed DNA gaps. Non-colocalised molecules in the *in-situ* formed gaps likely stem from non-specific binding of the Cy3B-modified flanking strand on the surface, or the absence of the Cy3B-containing flanking strand from the initially deposited DNA gap. Since the actual SPIN-seq experiment uses 10- to 20-fold higher concentration of flanking strands than in the characterization experiment here (with the latter being limited to 100 nM due to presence of a label on the flanking strand), we conclude that, during *in-situ* DNA gap formation using unlabelled flanking strands, >90% of DNAs is converted to gapped-DNA molecules. To verify the functionality of the *in-situ* formed DNA gap, we performed transient hybridization of the complementary seal on the *in-situ* formed DNA gap molecules and observed similar seal binding for the pre-formed and *in-situ* formed DNA gap molecules (**fig. S18D**).

To read position 25 using competitive inhibition, we used sequential interrogations with 200 nM of R-seal R9-CAP-N^5^-Cy5 and 600 nM of each of the 4 U-seals in IB3. In each round of interrogation, we collected 1200 frames using 250-ms exposures.

Preparation and single-molecule imaging of transcription complexes and reactions. Open complexes were formed by incubating 100 nM lacCONS DNA variant library with 150 nM of RNAP holoenzyme in KG7 buffer (40 mM HEPES pH 8, 100 mM potassium glutamate, 10 mM MgCl_2_, 1 mM DTT, 100 µg/mL BSA, 5% glycerol) at 37˚C for 30 min. Subsequently, 1 μL of 1 mg/mL heparin was incubated in the solution for 1 min, followed by centrifuging for 2 min at 2000 RPM. The supernatant was collected in a separate centrifuge tube, and 0.5 μL of the open complex mixture was incubated on a neutravidin-coated slide having 30 μL of KG7 as prepared for single-molecule DNA imaging. After washing off unbound open complex from the surface, we incubated the surface with 500 μM ApA dinucleotide (Jena Bioscience) diluted in KG7 buffer for 10 min followed by adding an oxygen scavenging system (1% glucose, 40 µg/ml catalase, and 0.1 mg/ml glucose oxidase). We started imaging the complexes using 532nm/638nm ALEX excitation and exposures of 200 ms; we then added 200 μM NTPs in real-time using an in-house made fluidic system equipped with magnetically controlled tubing positioner at ~15 s from starting imaging. We tracked the signal for 6.6 min, followed by washing the surface three times with 40 µL of MQ H_2_O.

In-situ DNA gap preparation and sequencing after transcription assay. After imaging transcription reactions, we washed the surface with MQ H_2_O, added 40 µL of 20 mM freshly prepared NaOH for 30 s to de-hybridize the dsDNA, and washed the surface three times with 40 µL of AB. To create an 8-nt gap around the sequence of interest (positions +6 and +7), two flanking sequences of 39-nt and 41-nt (lacCONS(+11/+49)-B and lacCONS(-39/+2)-B) were incubated on the surface for 15 min at 2 µM final concentration in IB3. Sequencing was performed using 200 nM of 8-nt R-seal and 8 sequential interrogations using a mixture of 200 nM R-seal (R8-lacCONS(+3/+9)-N^+7^N^+6^-B^A647N^) and 600 nM of each of the 8 U-seals (**table S1**) in IB4 (20 mM HEPES pH 7.7, 150 mM MgCl_2_, 200 mM NaCl, 10% dextran sulphate, 5% formamide and 200 µM of freshly prepared spermidine hydrochloride).

Image and time-series analysis. For all movies, drift correction was performed by first identifying localizations of the fiduciary markers (beads) on the surface using in-house python algorithm (<https://github.com/Z-Qing/GapSeq-repo>), followed by applying the AIM algorithm (*47*) to calculate the drift, and shifting the movies in the opposite direction to compensate for it. Acceptor and donor channels in the movies were aligned to the donor channel of the localization movie using the pystackreg library (*48*) with a rigid-body transformation. The transformation matrix between donor channels, as well as the transformation between the donor and acceptor channels, was computed using fiducial markers from the initial frames.

The aligned and undrifted movies were further analyzed using in-house software *napari-molseeq* (<https://github.com/piedrro/napari-molseeq>). First, gapped-DNA molecules were localized using the integrated Picasso algorithm (*49*) with a gradient of 500 and bounding box size of 5 pixels. For all gapped-DNA localizations, seal DNA binding traces were determined in a bounding box mask of 4 pixels and background mask of 1 pixel, keeping 1 pixel as a buffer zone between the bounding box and background mask. For the intensity determination, the sum and mean of all the 4 pixels inside the bounding box mask was used. Traces were exported in JSON format and visualized using in-house software *Traceanalyser* (<https://github.com/piedrro/TraceAnalyser>).

For all dwell-time analysis, traces for each experiment were fitted using a 2-state Hidden Markov model (integrated in *traceanalyser*) for 1-base sequencing and an automatic state selection model for 3-base and 5-base sequencing. Histograms and traces were plotted in OriginPro 2021 (OriginLab). Background subtraction was carried out using a median filter with a window size of 1 pixel. The resulting images were scaled and translated to bring them into the same intensity range before performing the registration.

Base calling. For base calling on each gapped-DNA molecule (identified in the green channel), we used in-house Python-based tools (<https://github.com/Z-Qing/GapSeq-repo>) to count the number of localizations due to the binding of labelled seal-DNA in the red channel. Localizations in the green channel were determined using Picasso (*46*) with a gradient threshold of 400. Localizations in the red channel were found using a gradient threshold of 1000, with the following camera parameters: baseline = 400, box size = 5, EM gain = 1, quantum efficiency = 0.82, sensitivity = 2.5, and pixel size = 117 nm. We used the same parameters for all movies collected throughout all the experiments.

For each localization in the donor channel, the number of localizations in all acceptor channels within a radius of 2 pixels was counted. To define a threshold number of localizations above which only we will call the bases (a “base-calling threshold”), we first counted all localizations from all the interrogation movies for one position and acquired the probability density function (PDF) of the localisation counts using a Gaussian kernel (**fig. S5A**). An exponential function was then fitted to the distribution of first derivative of the PDF, starting from its global minimum (**fig. S5B**). The final base-calling threshold was algorithmically set to the value on the x-axis corresponding to this global minimum, plus three times the decay-length parameter obtained from the exponential fit.

Subsequently, we performed additional filtering to remove molecules irreversibly adsorbed to the surface near the gapped DNA, but not interacting with DNA; such molecules show persistent localisation clustering over time. To remove such molecules (that show at least 100 localisations), we first assigned localisations in consecutive frames to a “localisation bundle”. There should be at least 5 bundles (for single-base calling) or 2 bundles (for sequencing multiple bases), and the longest bundle should be smaller that 50% (for single-base calling) or 80% (for sequencing multiple bases) of the frame number. When both conditions are met, the molecules are accepted, otherwise rejected.

For every gap localization, we found the four localization counts for each of the seal-interrogations and took the difference between the largest and second largest localization counts. Confidence of calling the base is defined as following:

$${difference}_{non-comp}=largest count-second largest count$$

$${confidence}_{non-comp}=\frac{difference}{(95 percentile of difference)}$$

This logarithmic transformation enhances the dynamic range and stability of the confidence metric. Selecting the gap molecules with higher confidence increase the accuracy of sequencing (to ~98%) while still maintains most molecules (**fig. S5C**).

In the competitive-inhibition approach, 3 out of 4 traces exhibit more prominent binding. Using a similar base-calling threshold, we chose the trace exhibiting the least number of localizations as the one complementary to the sequence at the corresponding gap position. To determine a confidence metric for base-calling, we used the difference between the second-lowest localization count and the lowest localization count values out of the four U-seal interrogations.

$${difference}_{comp}=second least count-least count$$

$${confidence}_{comp}=\frac{difference}{(95 percentile of difference)}$$

Data analysis for CAP-binding assay and *in-situ* sequencing. All movies were undrifted and aligned, and traces were determined as described above. Traces were then fitted using HMM model (automatic state selection). All traces exhibiting binding were exported in CSV format. Traces were then fitted using a cluster-based approach for dwell-time analysis (<https://github.com/Z-Qing/GapSeq-repo/tree/main/trace_analysis>). Base calling for the localized CAP-binding DNA molecules was done using the base-calling algorithm in the previous section. Bases are called above a confidence threshold value of 0.4.

Dwell times and identified bases were aligned using in-house MATLAB script. Dwell-time and unbound-time histograms were fitted using double-exponential and mono-exponential decay functions, respectively, in Origin (OriginPro2022). We then determined the rates of binding, unbinding, and the equilibrium binding constant using following equations:

$$k_{off}=\frac{1}{mean dwell time}$$

$$k_{on}= \frac{1}{concentration of CAP\times mean unbound time}$$

$$K_{d}=\frac{k_{off}}{k_{on}}$$

Data analysis for transcription assay and *in-situ* sequencing**.** All movies were undrifted and aligned, and traces were determined as described above. For all further trace analysis and sequence calling, we used localizations in the green channel (which is also the donor channel for the smFRET measurements) collected immediately after the immobilization of promoter library. All sequencing movies were aligned using the fiduciary marker positions in the green channel during the transcription assay movie. Using a 4-pixel mask and 1-pixel background mask, we determined intensity vs time trajectories of the intensity of donor emission upon donor excitation (*I_DD_*), the intensity of acceptor emission upon donor excitation (*I_DA_*), and the intensity of acceptor emission upon acceptor excitation (*I_AA_*). Traces obtained for transcription assay were further visualized and analyzed using ‘Traceanalysis’ software. Intensity vs. time trajectories were curated to exclude multiple step donor and acceptor photobleaching, and trajectories exhibiting donor and acceptor photoblinking before or during E_FRET_ reaching 0.4.

Apparent FRET efficiency (E_FRET_) was determined using the following formula

$$E_{FRET}= \frac{I_{DA}}{{(I}_{DA}+I_{DD})}$$

Intensity time-traces exhibited transitions between various distinct E_FRET_ states possessing anti-correlated *I_DD_* and *I_DA_* changes. FRET efficiency time-traces were fitted using multi-state HMM model as implemented in *Traceanalysis* software. Molecules were then classified to Classes I to IV based on fluctuations between FRET states (see main text).

Base-calling was performed using the competition-assay base-calling algorithm described earlier, and sequences were linked to FRET time-traces. We excluded FRET time-traces for which the base calling was not possible due to the localizations count being below the basic-calling threshold, or due to U-seal interrogations exhibiting a similar localization count for all four.

For each time-trace with an assigned sequence, the pause time was determined using the dwell-time at E_FRET_~0.4 (corresponding to the paused complex at ~6-nt RNA) for Classes I, II and III. For Class II molecules, we considered only the first E_FRET_~0.4 dwell. Pause-time histograms (**Fig. 5F**, **fig. S24**) were plotted, binned, and fitted using mono-exponential decays in Origin2021. Propensities for each class of molecules for different sequences were determined by dividing the molecules falling into each class by the total number of transcribing sequenced molecules for that sequence. Class-propensity heatmaps (**Fig. 5F-I**) were plotted in Origin2021.

**Supplemental Figures**

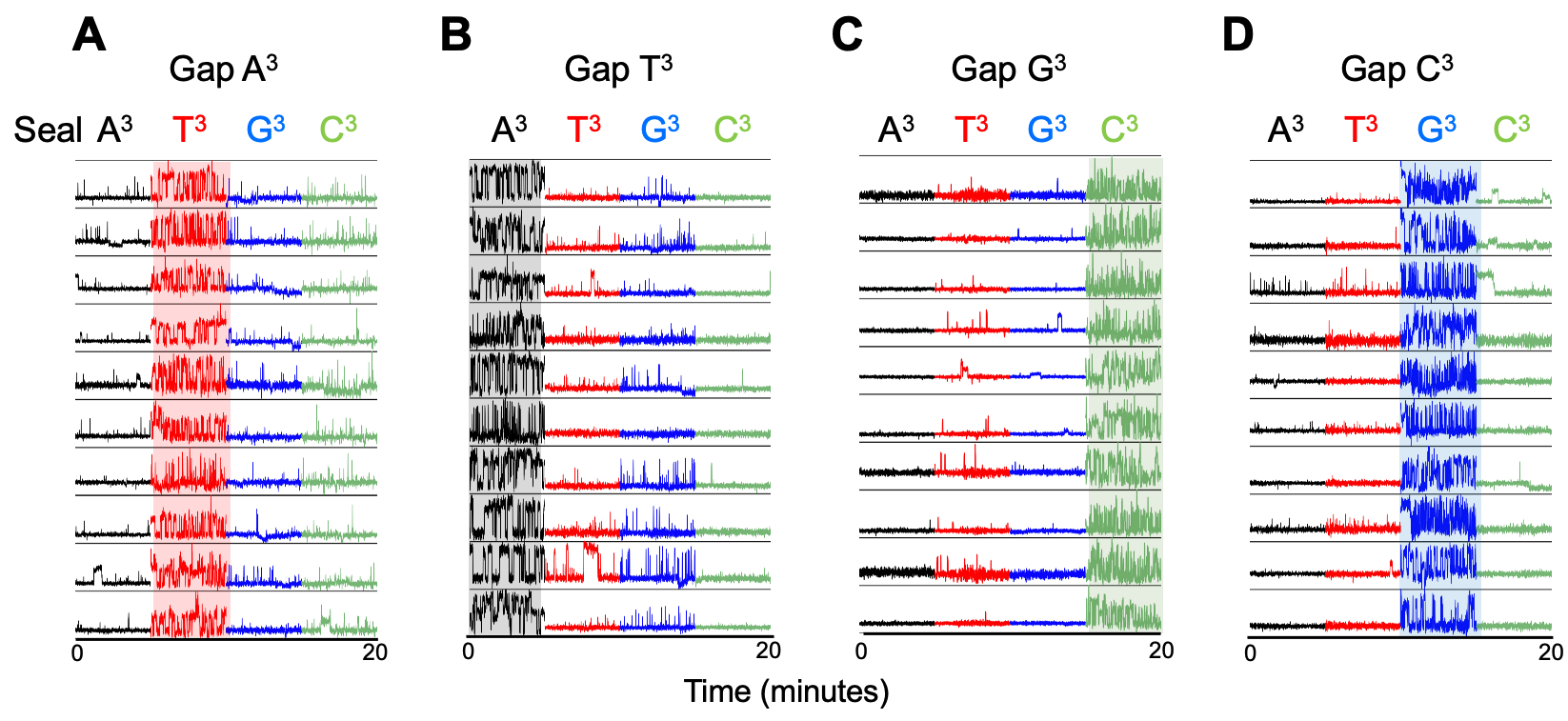

Fig. S1. Additional example traces for identification of single nucleotide differences using transient DNA hybridization.

**A**. An 8-nt DNA gap is sequentially interrogated at gap position 3 (where the base is an A) using 4 seals that differ only on the base opposed to the interrogated position. Only the seal featuring A at position 3 (red traces) shows strong gap binding; the remaining seals show negligible binding.

**B-C.** Analysis as in panel A, but for gaps featuring a T, G, and C at gap position 3.

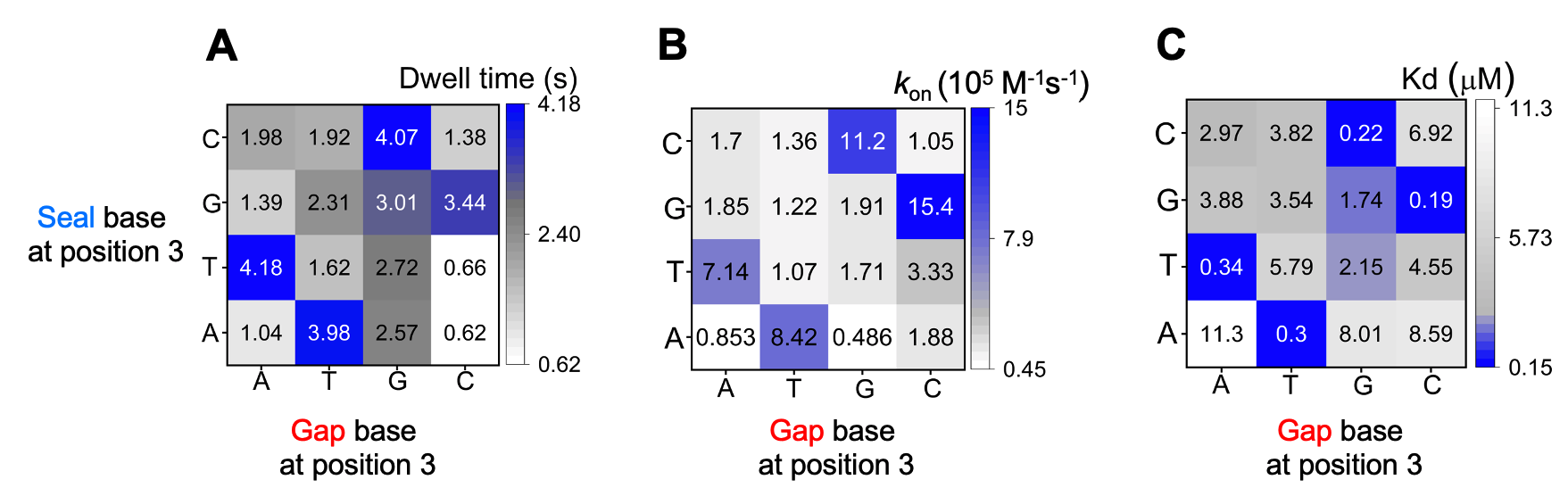

**Fig. S2. Kinetic and thermodynamic analysis of interactions of complementary and 1-nt mismatched seals with gapped-DNA.**

**A.** Dwell times of different 8-nt seals (each featuring a different standard base corresponding to gap position 3) on 8-nt gaps (each featuring a different standard base at gap position 3). Fully complementary pairs exhibit longer dwell times compared to 1-nt mismatched pairs.

**B.** Binding on-rates for the same pairs of interactions in panel A. Fully complementary pairs exhibit faster on-rates compared to 1-nt mismatched pairs.

**C.** The dwell times and on-rates were used to create a similar heat map for equilibrium dissociation constants (K_d_ values). Fully complementary pairs exhibit lower K_d_ (higher affinity) compared to 1-nt mismatched pairs.

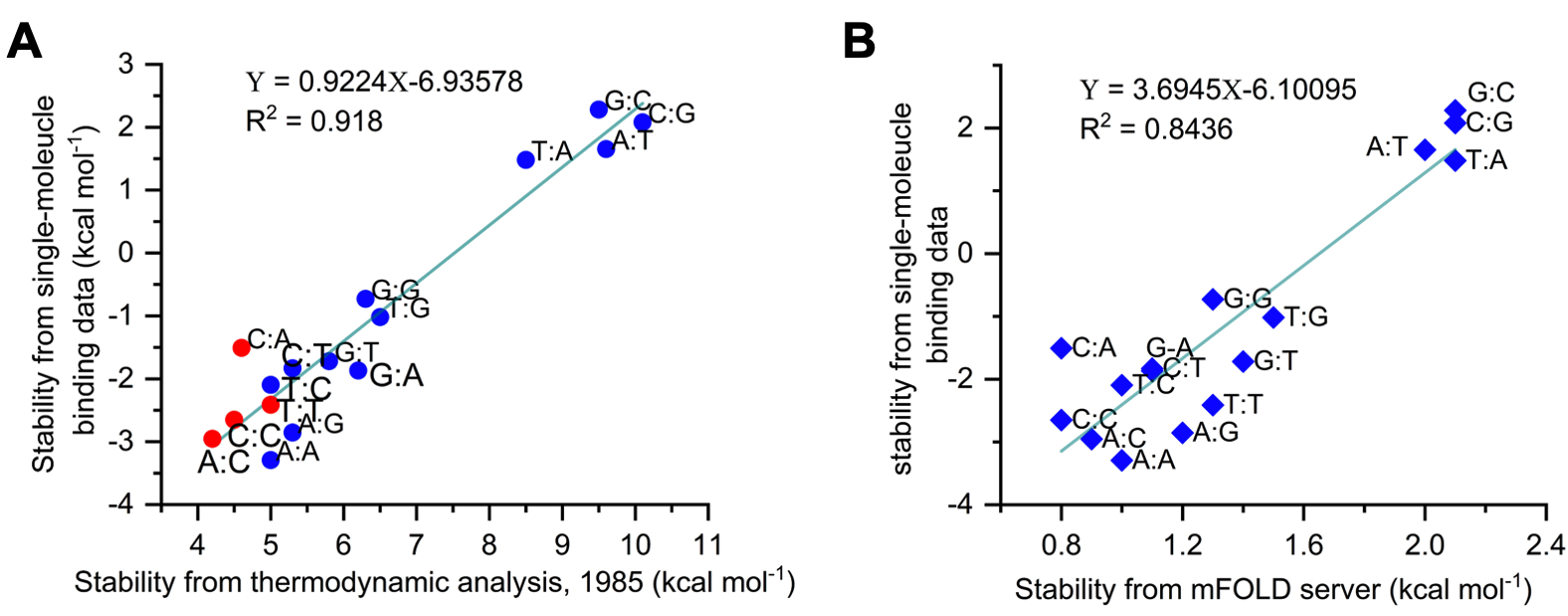

**Fig. S3. Comparison of thermodynamic stability of transient gap-seal duplexes as measured at the single-molecule level with previous analysis using experimental and computational approaches.**

**A.** Comparison of stabilities obtained from single-molecule analysis (fig. S2) with the stability obtained from spectrophotometric thermodynamic analysis *(15)* which looked at the mismatches of base X in an A-X-T context; our sequence context is C-X-A. The results from both analyses are highly correlated (R^2^ = 0.918). Red points are associated with higher uncertainty in the ensemble spectrophotometric analysis.

**B.** Comparison of stabilities obtained from single-molecule analysis (fig. S2) with stability estimates the DINAmelt server *(17)* using the context used in our experiments. An additional base-pair was added to the 8-mer to emulate the effect of stacking provided by the two ends of the gap. The results are highly correlated (R^2^ = 0.844).

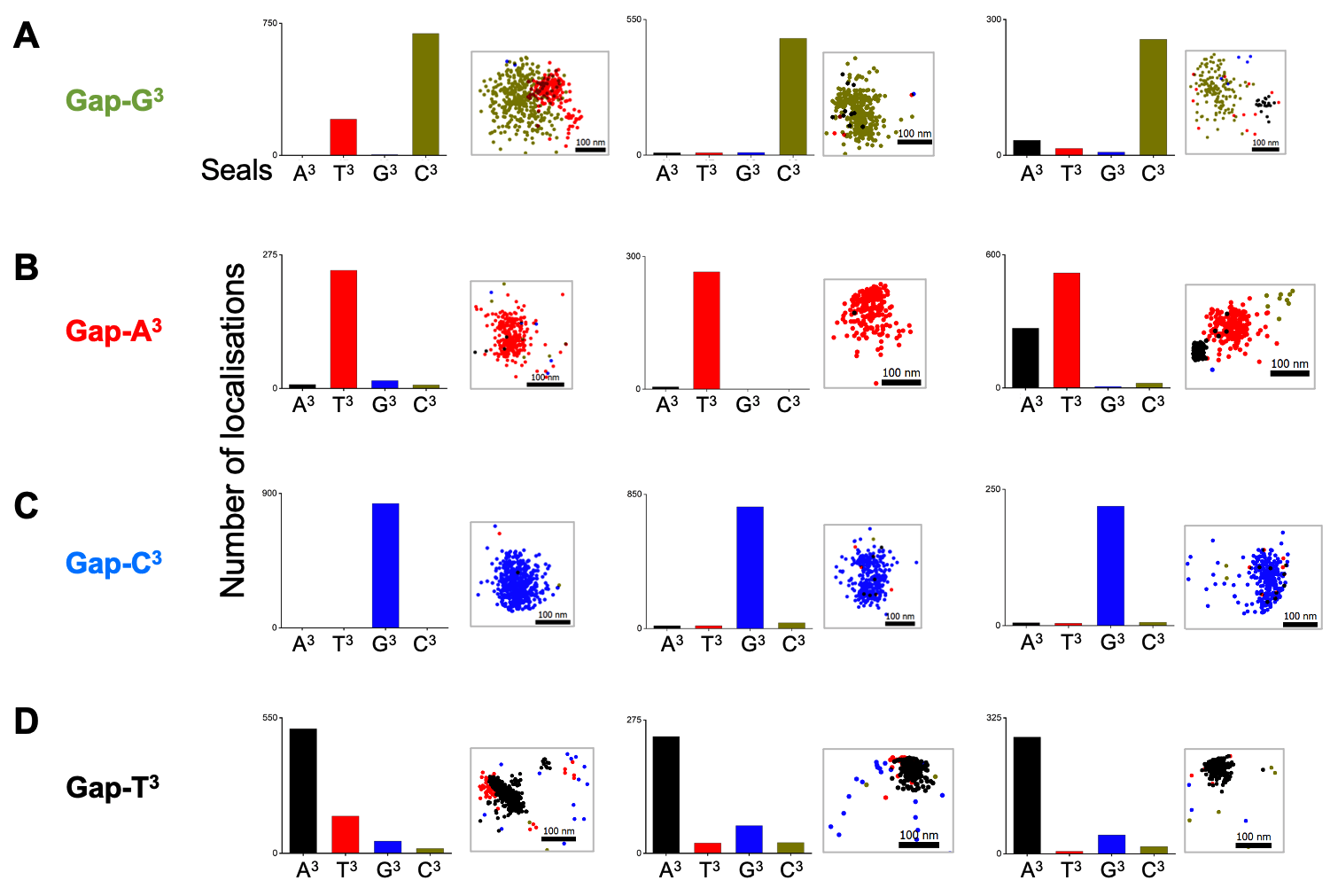

**Fig. S4. Quantification of seal binding using localization counting.** Representative plots for single-molecule localization counts and locations for seal binding to gap molecules featuring a different base at gap position 3 (Gap-G^3^, Gap-A^3^, Gap-C^3^, and Gap-T^3^).

**A.** Bar plots: single-molecule localisation count for three example Gap-G^3^ molecules sequentially interrogated with 4 labelled seals; in all three cases, we observe many more localizations for the complementary seal (seal C^3^). Scatter plots: maps of seal-binding localizations; localisations due to the complementary seal are clustered in near-circular pattern; occasionally, mismatch seals bind to a proximal (but different) location (e.g., in the first example, seal T^3^ binds ~100 nm away from the main cluster), leading to background; these results point to the possibility of using spatial filtering to improve base-calling accuracy.

**B-D.** As in A, but for gap molecules Gap-A^3^, Gap-C^3^, and Gap-T^3^.

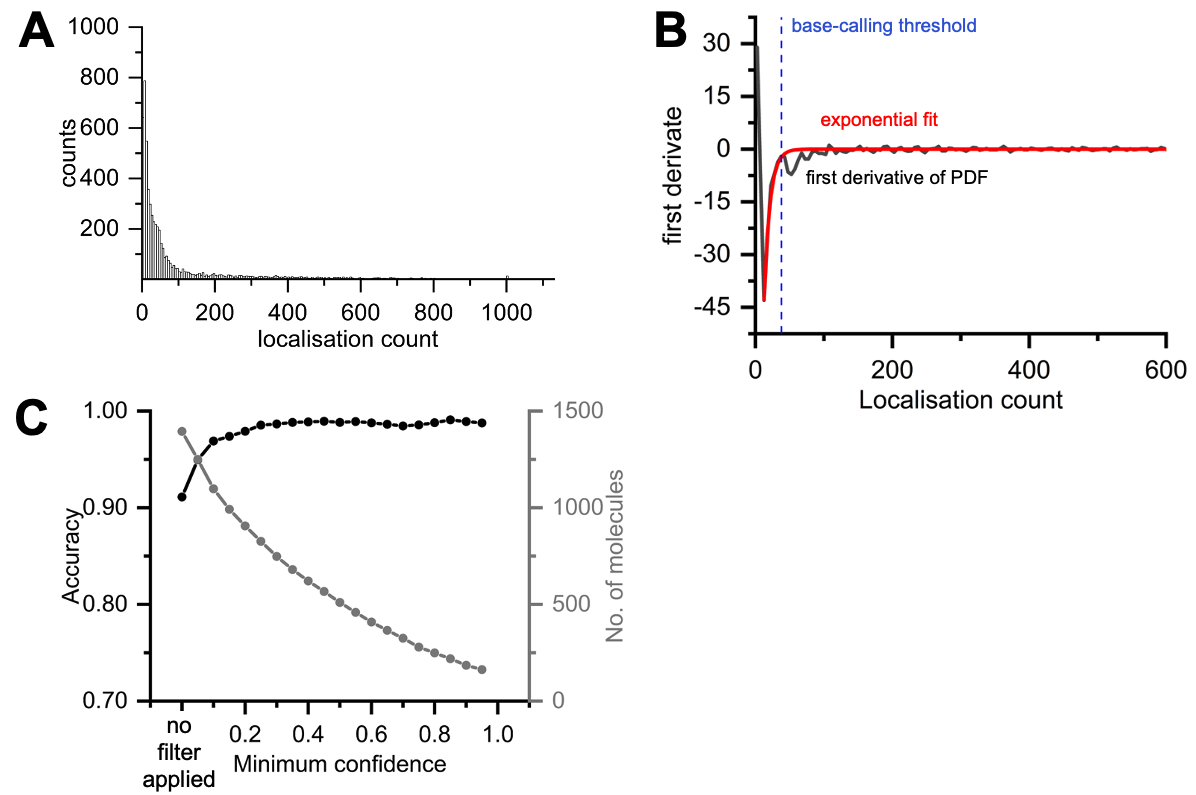

**Fig. S5. Base-calling method and accuracy test.**

**A.** Frequency histogram of all localization counts due to the four seals binding to Gap-G^3^.

**B.** First derivative of the probability density function of the frequency of localisation counts (black line) and fit to an exponential decay function (red line). Blue line, threshold value of localization count; the threshold is defined as the localisation count of first derivative minimum (in this example, 13 localisations) plus three times the decay length (in this example, 8 localisations) obtained from fitting. So, the final base-calling threshold in this example is set to 37 localisations. After filtering, we keep only molecules for which at least one localisation count (out of four interrogations) is higher than the threshold.

**C.** Accuracy and number of correctly called molecules with increasing confidence value (see SI text for the definition of confidence values).

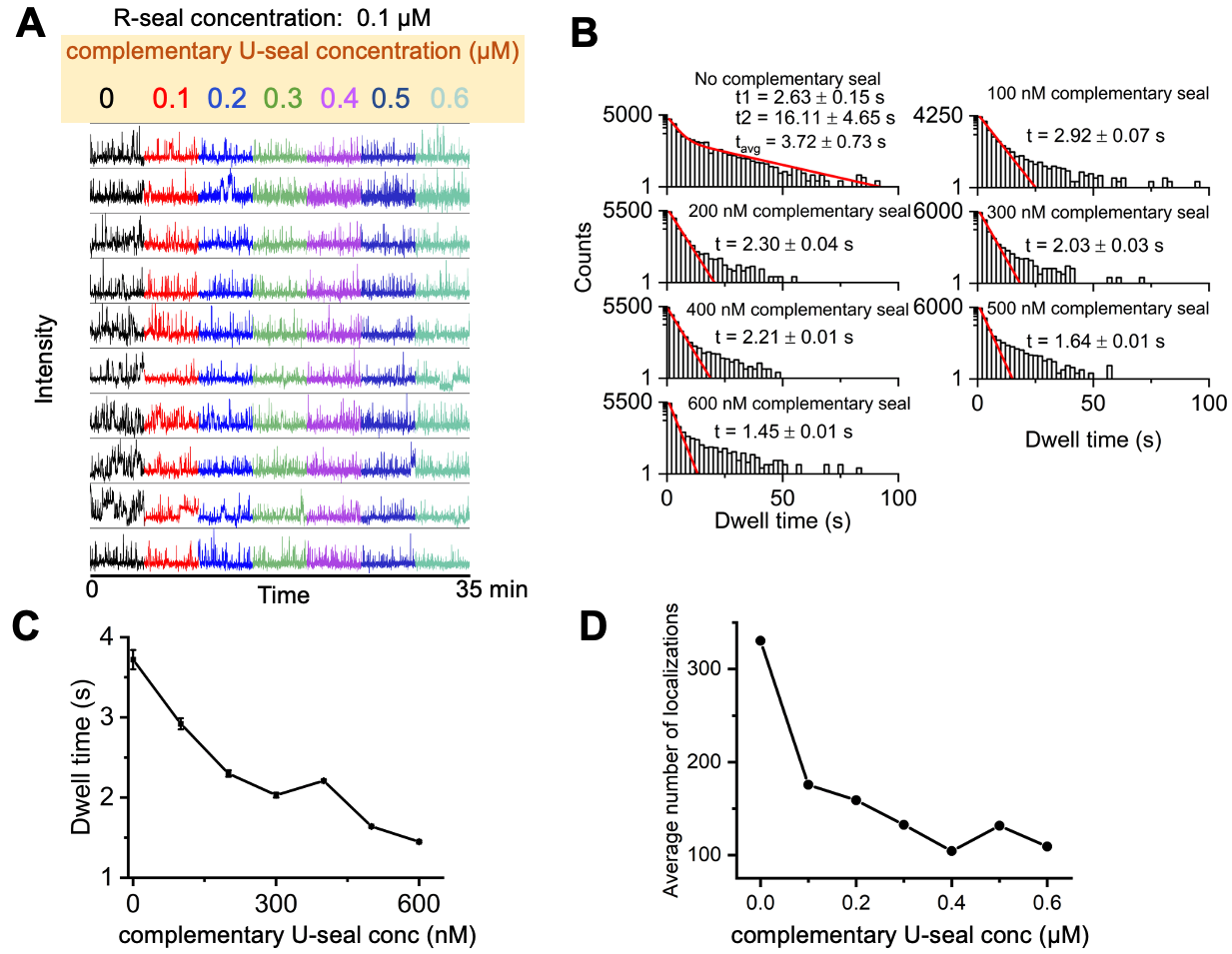

**Fig. S6. Quantifying the effect of competitive inhibition of R-seal binding to a gap due to increasing concentrations of a complementary U-seal.**

**A.** Representative traces for gap-binding of 100 nM of R-seal R8-N^3^ in presence of 100-600 nM complementary U-seal in IB2 buffer.

**B.** Dwell time distributions for the different U-seal concentration. Histograms are fitted with double-exponential decay model (red lines).

**C-D.** Mean dwell-times (panel C) and localisation count (panel D) as a function of U-seal concentration.

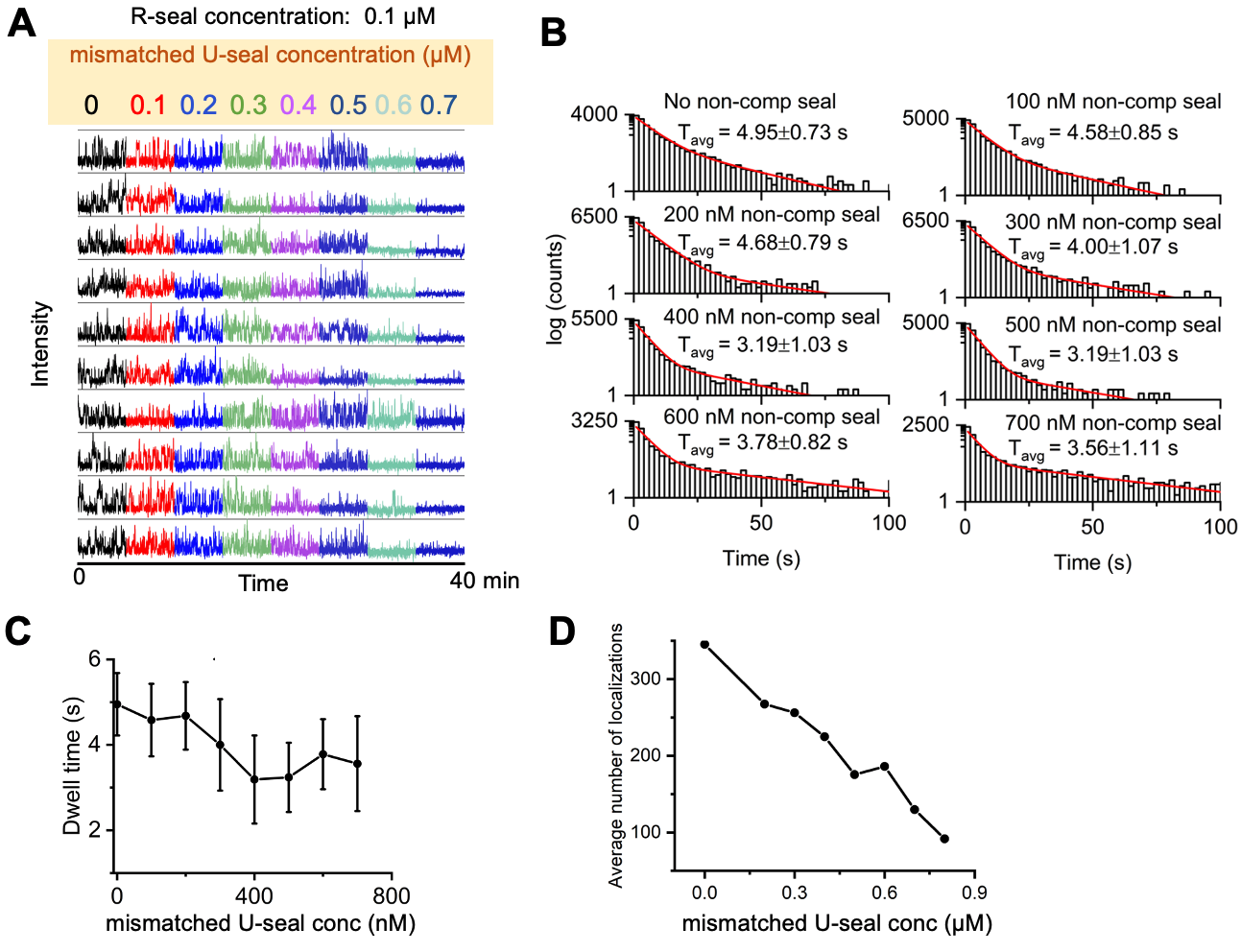

**Fig. S7. Quantifying the effect of competitive inhibition of R-seal binding to a gap due to increasing concentrations of a 1-nt mismatched U-seal.**

**A.** Representative traces for gap-binding of 100 nM of R-seal R8-N^3^ in presence of 100-600 nM 1-nt mismatched U-seal (seal U8-T^3^) in IB2 buffer.

**B.** Dwell time distributions for the different U-seal concentration. Histograms are fitted with double-exponential decay model (red lines).

**C-D.** Mean dwell-times (panel C) and mean localisation count (panel D) as a function of U-seal concentration.

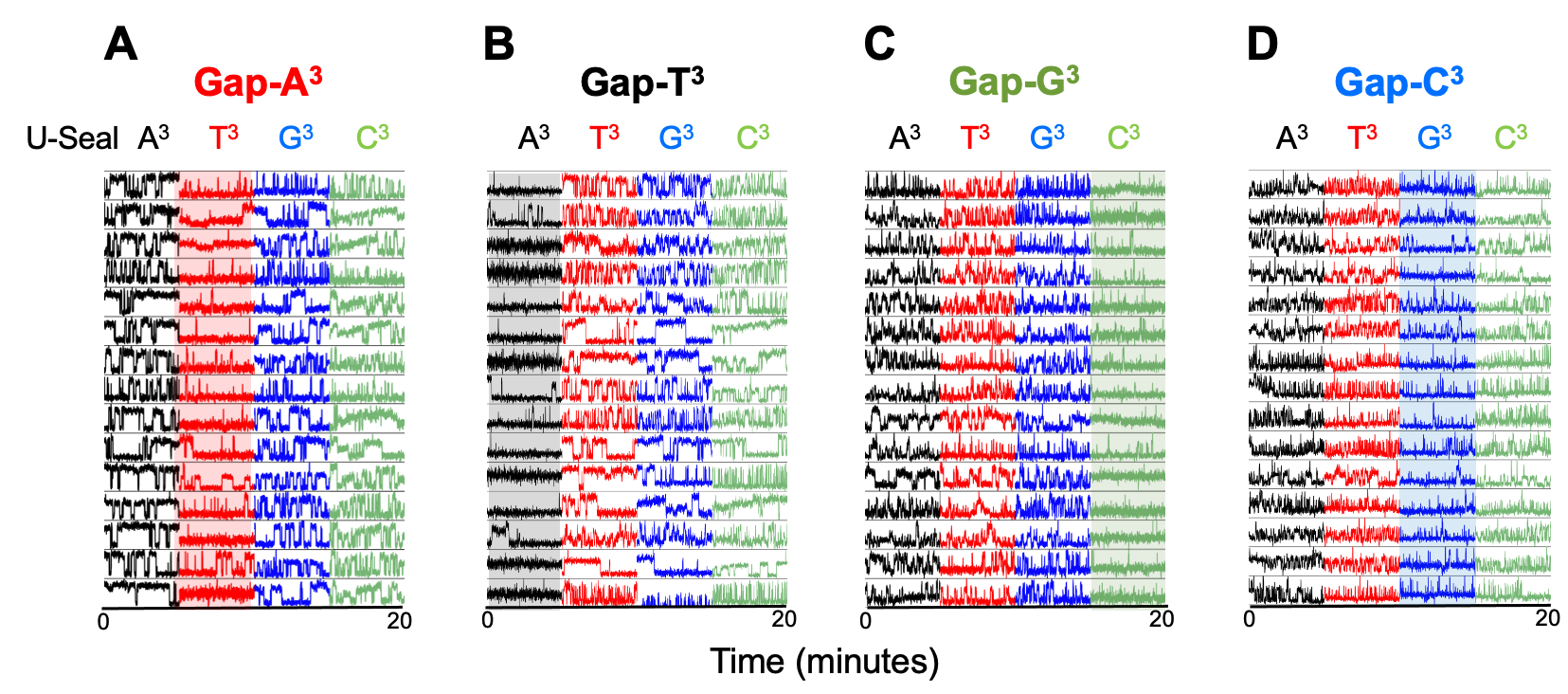

**Fig. S8. Reading single nucleotides using competitive inhibition: additional examples**.

**A.** Examining gap position 3 in Gap-A^3^ using 200 nM of labelled R-seal and a set of 4 unlabelled U-seals (added sequentially, all at 600 nM); the complementary U-seal (U8-T^3^, in red) suppress R-seal binding, enabling base identification.

**B-D**. Same analysis, but for gapped DNA molecules Gap-T^3^, Gap-G^3^, and Gap-C^3^. Complementary U-seals lower the extent of R-seal binding (see highlighted columns of timetraces), enabling base identification.

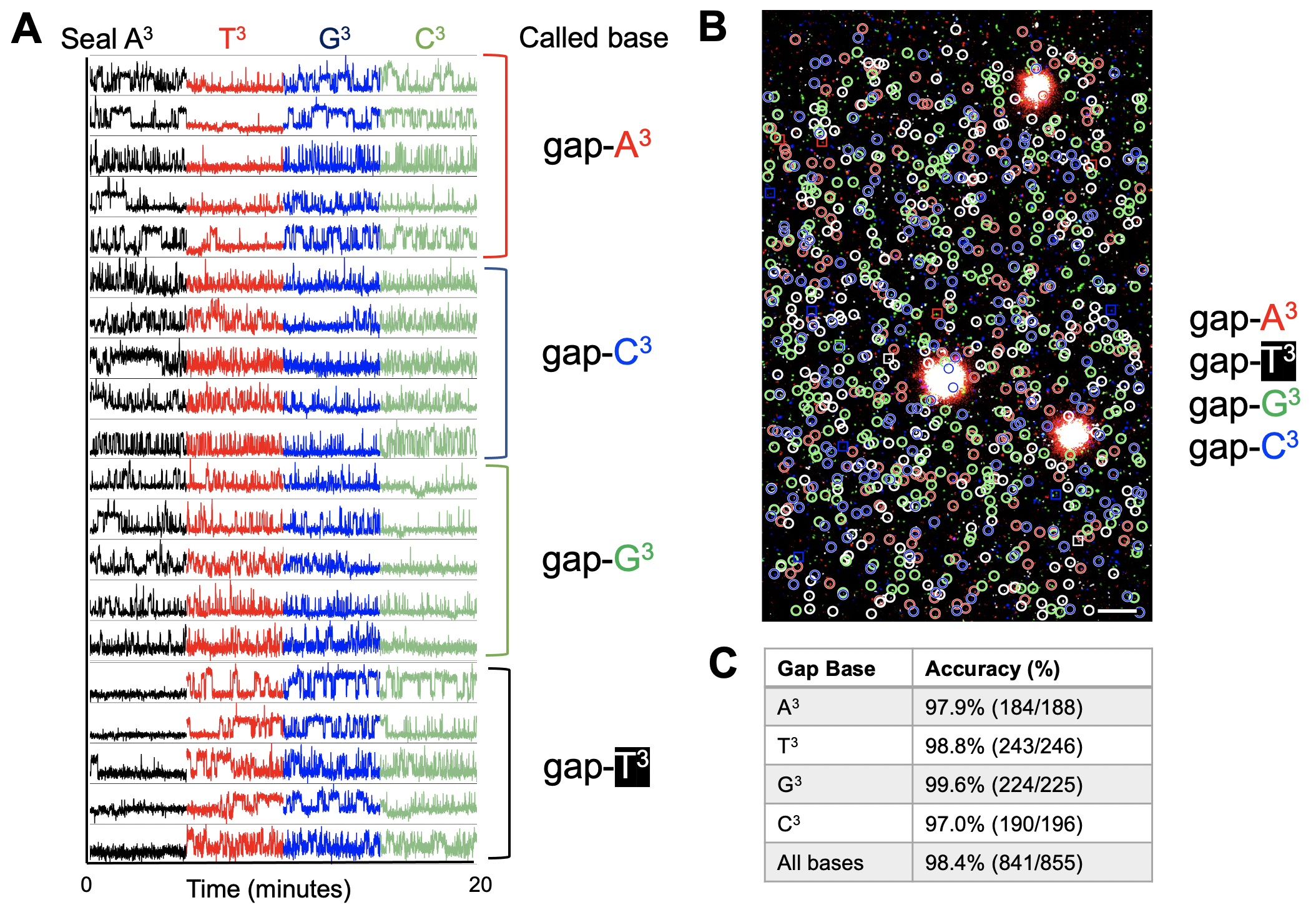

**Fig. S9. Validation of the competitive-inhibition approach using a mixed population of sequences with prior information.** All four gaps were deposited sequentially (as in **Fig. 2E**), followed by registration of their location on the surface, and photobleaching. DNA gaps were then interrogated using competitive inhibition using 200 nM of 8 nt R-seal and four cycles of interrogation with competing U-seals, each at 600 nM.

**A.** Sequential interrogation of the deposited DNA by seals specific for gap position 3. The absence of binding identifies the base in the interrogated position.

**B.** Field of view (50x80 μm) of molecules sequenced for gap position 3. White spots: fiducial markers. Circled molecules were identified correctly; in-square molecules were assigned incorrectly. The base-calling accuracy was ~98.4%. Scale bar: 5 µm.

**C.** Base-calling accuracies for individual bases.

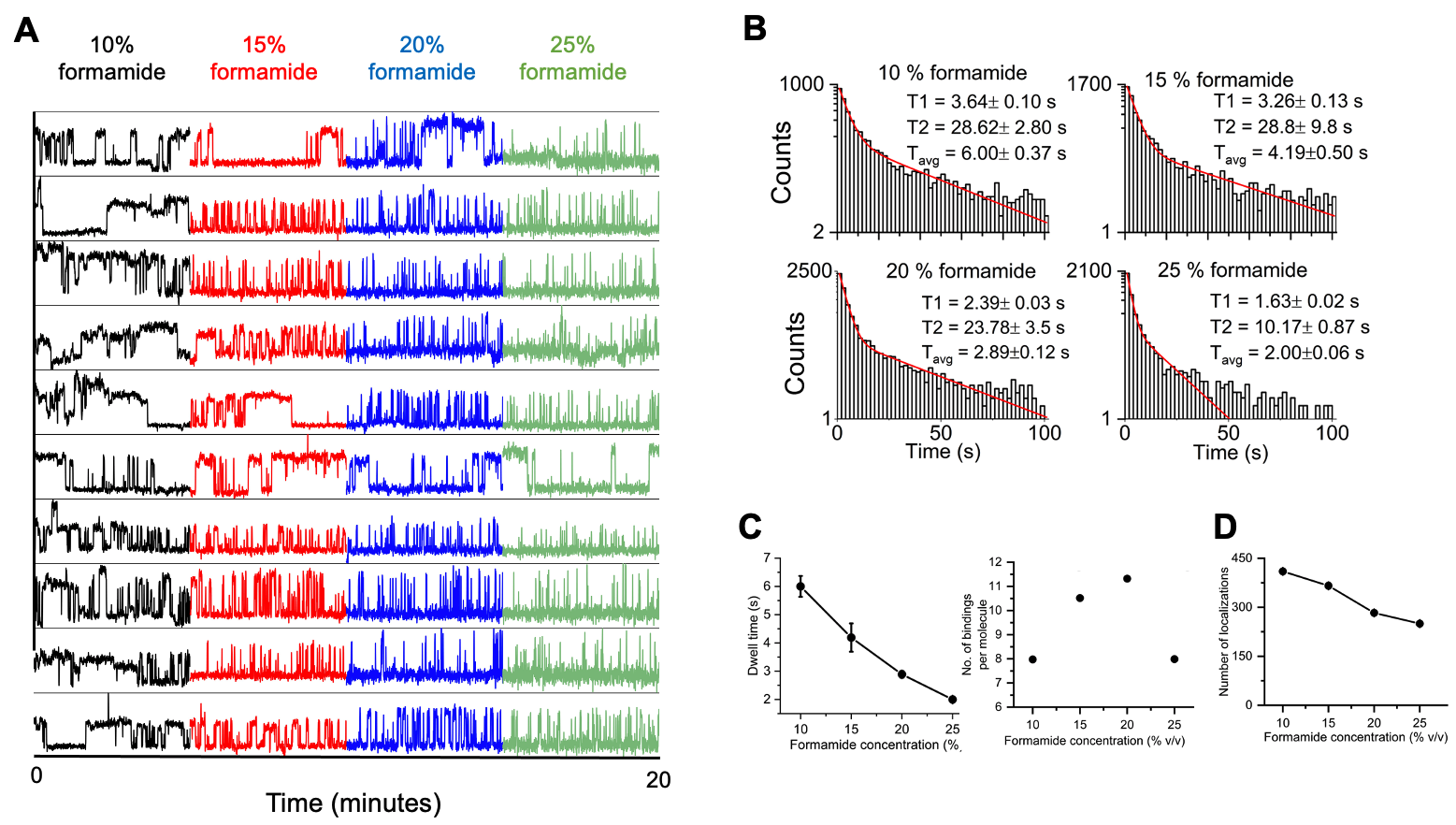

**Fig. S10. Optimizing formamide concentration for 13-nt seal binding of a gap.**

**A.** Representative traces from hybridization of 500 nM R-seal in IB3 (which contains 10% formamide) and in the presence of 10-25% formamide. Increasing formamide shortens dwell times in the bound (high-intensity) state.

**B.** Dwell-time histograms for the bound states at different formamide concentrations. Histograms are fitted with double-exponential decay model (red lines).

**C.** Mean dwell times in the bound states (top) and number of binding-unbinding cycles per gap molecule (bottom) as a function of formamide concentration. Formamide concentrations of 15% and 20% provide an optimal combination of mean dwell time and number of binding-unbinding cycles.

**D.** Mean localisation count as a function of formamide concentration.

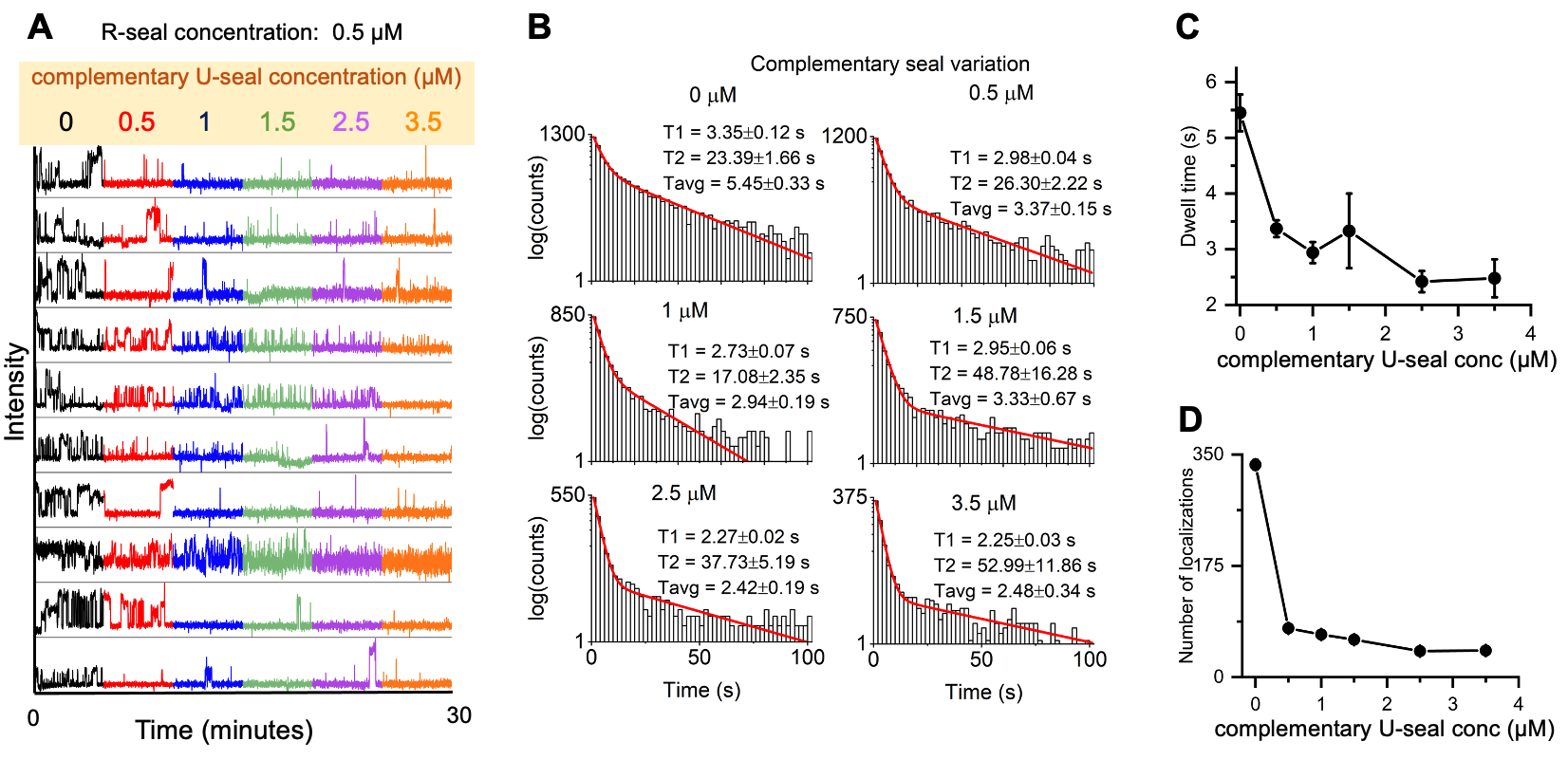

**Fig. S11. Quantifying competitive inhibition of R-seal binding to a gap due to increasing concentrations of a complementary U-seal: 3-base sequencing.**

**A.** Representative traces for gap-binding of 0.5 μM of R-seal R13-N^5^N^6^N^7^ in presence of 0.5-3.5 μM complementary U-seal (U13-C^5^N^6^N^7^).

**B.** Dwell time distributions for the different U-seal concentrations. Histograms are fitted with double-exponential decay model (red lines).

**C-D.** Mean dwell-times (panel C) and mean localisation count (panel D) as a function of U-seal concentration.

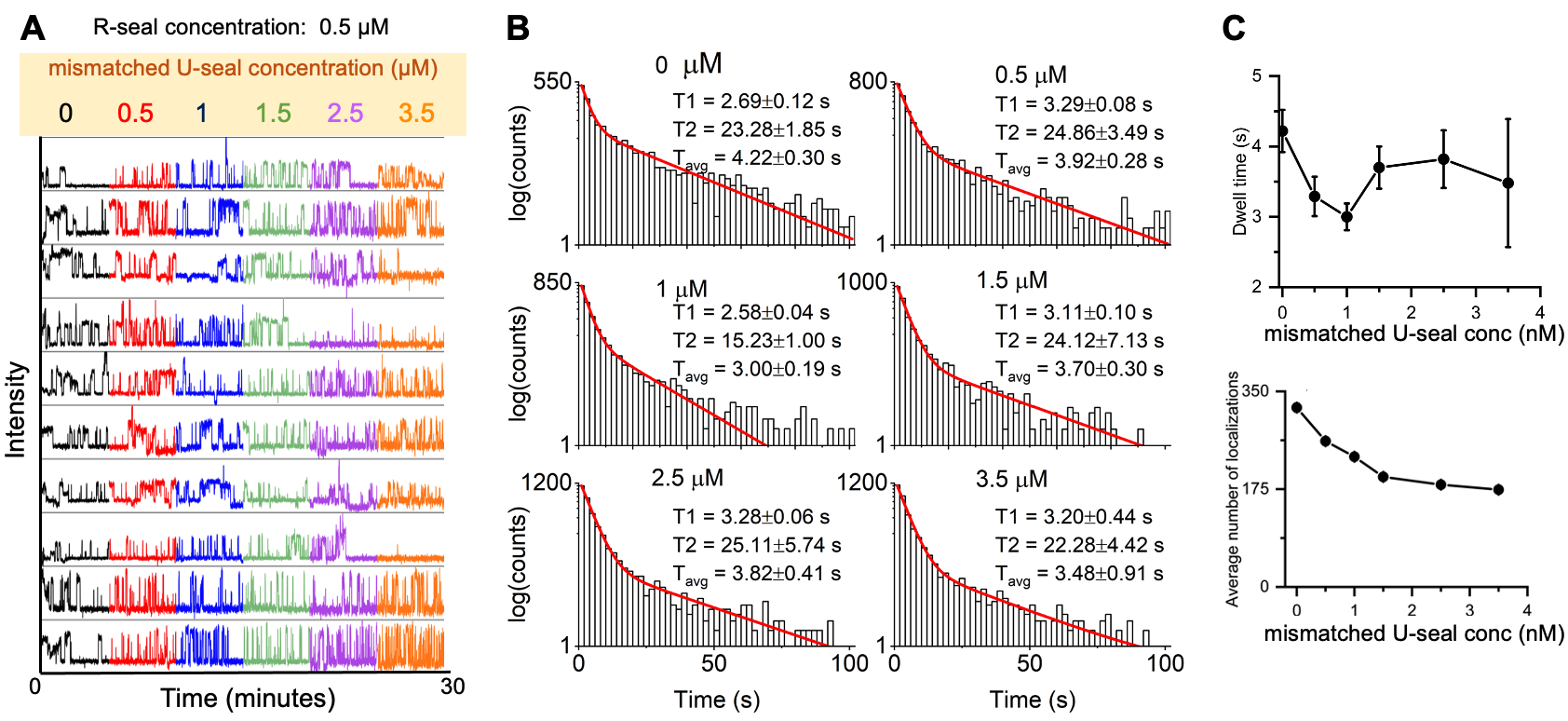

**Fig. S12. Quantifying the effect of competitive inhibition of R-seal binding to a gap due to increasing concentrations of a 1-nt mismatched U-seal: 3-base sequencing.**

**A.** Representative traces for gap-binding of 0.5 μM of R-seal R13-N^5^N^6^N^7^ in presence of 0.5-3.5 μM 1-nt mismatched U-seal (seal U13-T^5^N^6^N^7^ U13-T^3^).

**B.** Dwell time distributions for the different U-seal concentration. Histograms are fitted with double-exponential decay model (red lines).

**C-D.** Mean dwell-times (panel C) and localisation count (panel D) as a function of U-seal concentration.

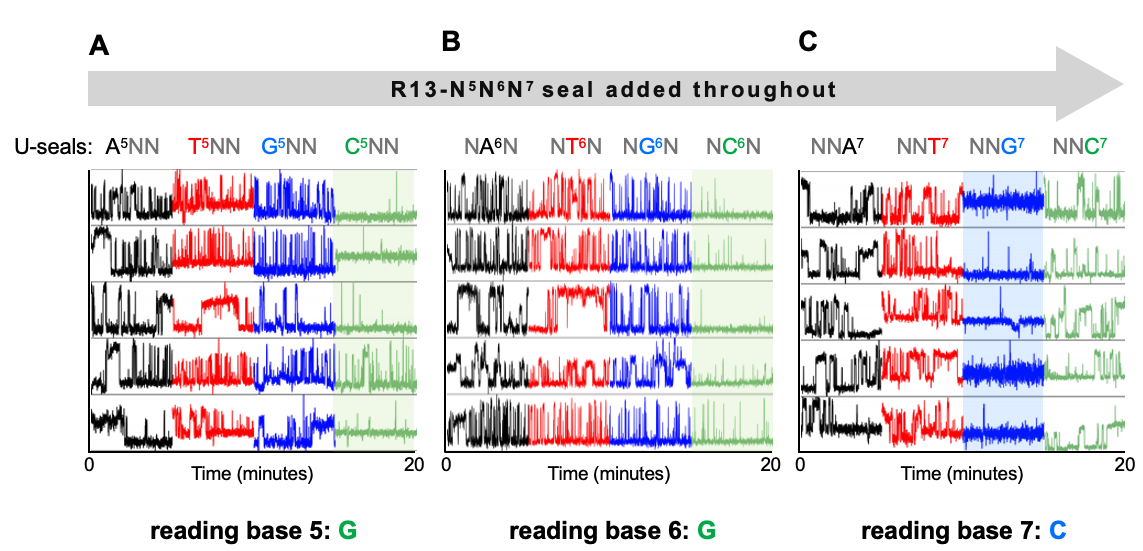

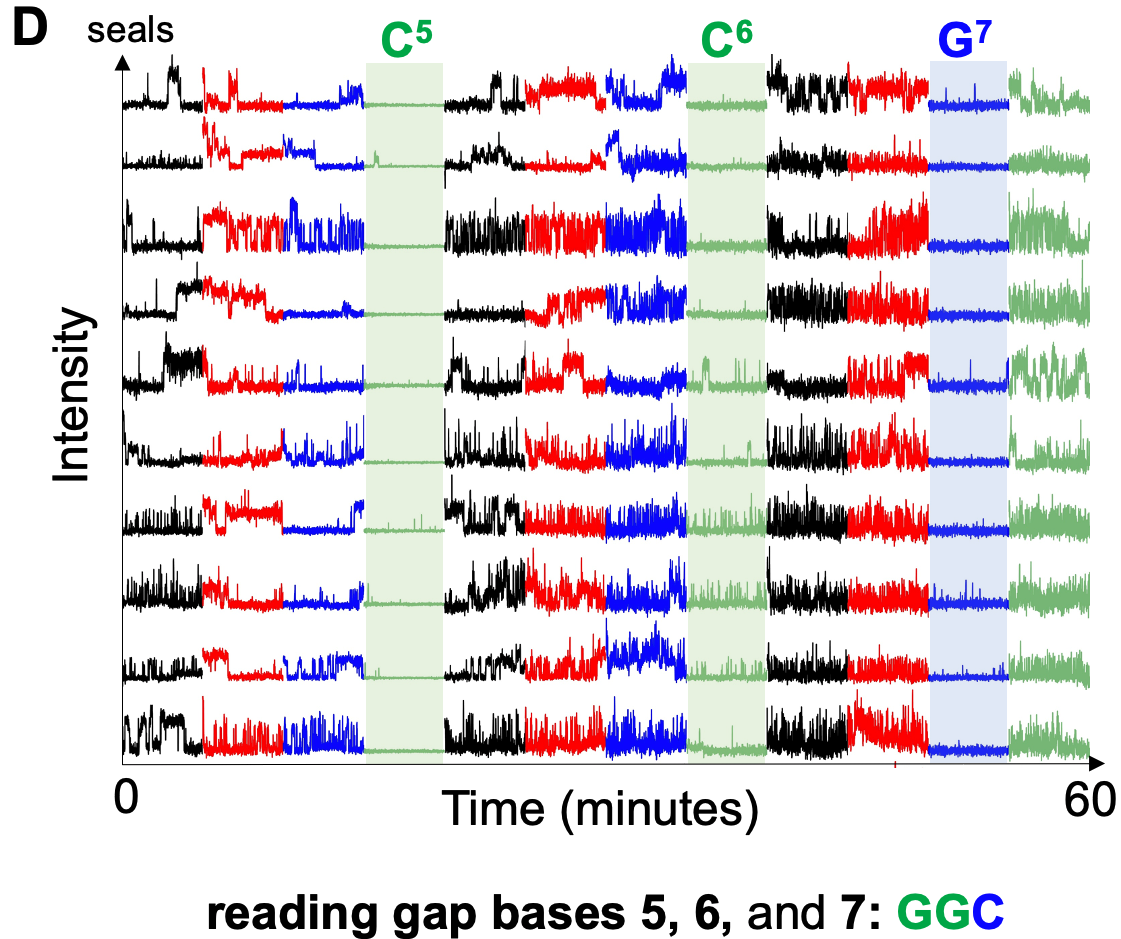

**Fig. S13. Reading three nucleotides using competitive inhibition**. **A-C.** Additional example traces for 3-base sequencing are provided for gap positions 5 (panel **A**), 6 (panel **B**), and 7 (panel **C**). Conditions and descriptions as in **Fig. 3F**. In all cases, the R-seal (R13-N^5^N^6^N^7^) was kept at 500 nM, and each base was evaluated on a different surface. Interrogation with the complementary U-seal led to decrease binding (highlighted traces).

**D.** Sequential interrogation of the same sequence (GGC) on DNA molecules immobilized on the same surface. The overall accuracy for molecules that give reads for all three bases is ~99.5% (188/189), whereas the accuracy for the individual positions is 97.8% (393/402), 91.6% (318/347), and 96.7% (911/942) for bases 5, 6, and 7, respectively.

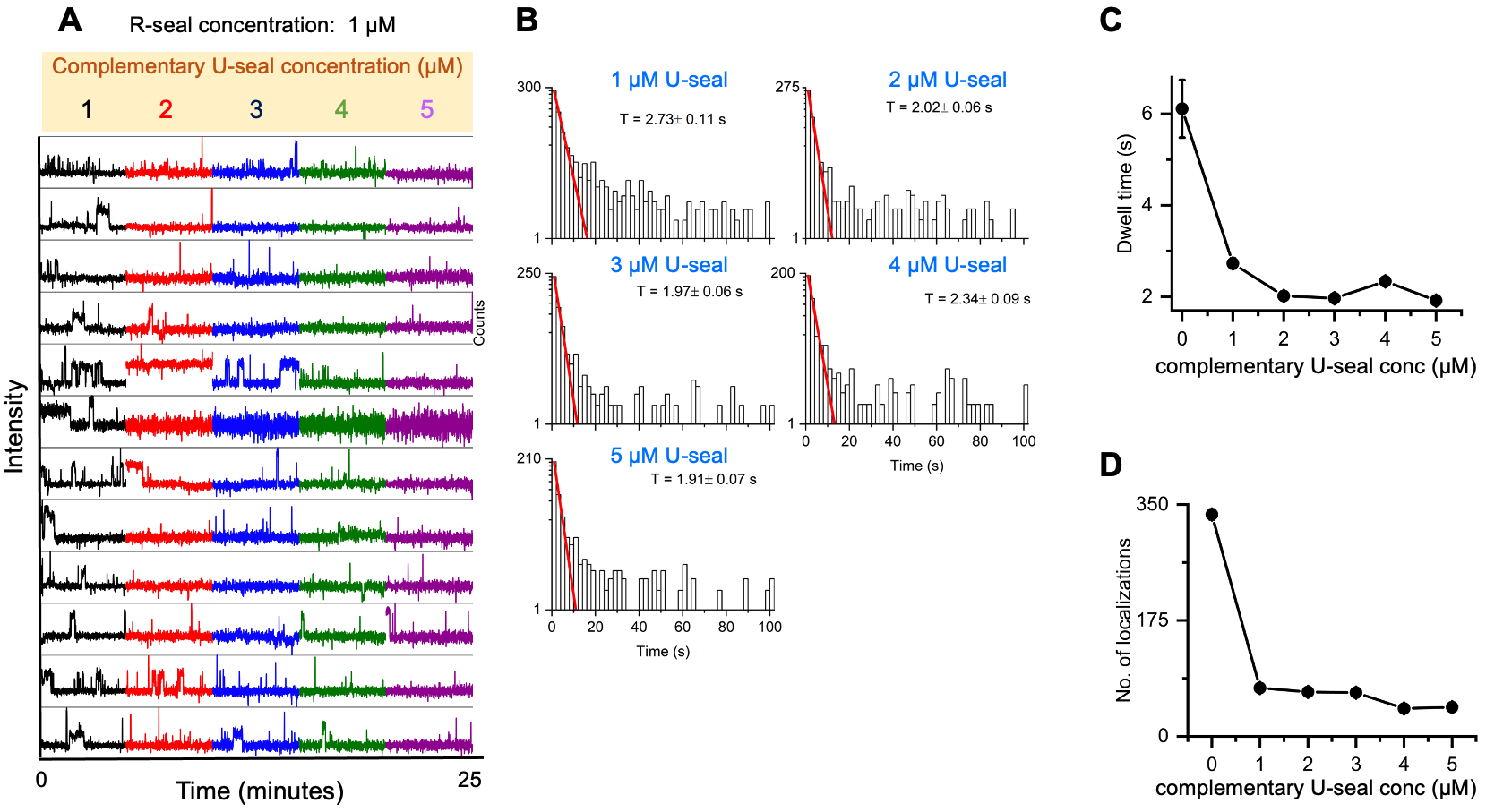

**Fig. S14. Quantifying competitive inhibition of R-seal binding to a gap due to increasing concentrations of a complementary U-seal: 5-base sequencing.**

**A.** Representative traces for gap-binding of 1 μM of R-seal R13-N^5^N^6^N^7^N^8^N^9^ in presence of 1-5 μM complementary U-seal (U13-C^5^N^6^N^7^N^8^N^9^).

**B.** Dwell time distributions for the different U-seal concentrations. Histograms are fitted with mono-exponential decay model (red lines).

**C-D.** Mean dwell-times (panel C) and mean localisation count (panel D) as a function of U-seal concentration.

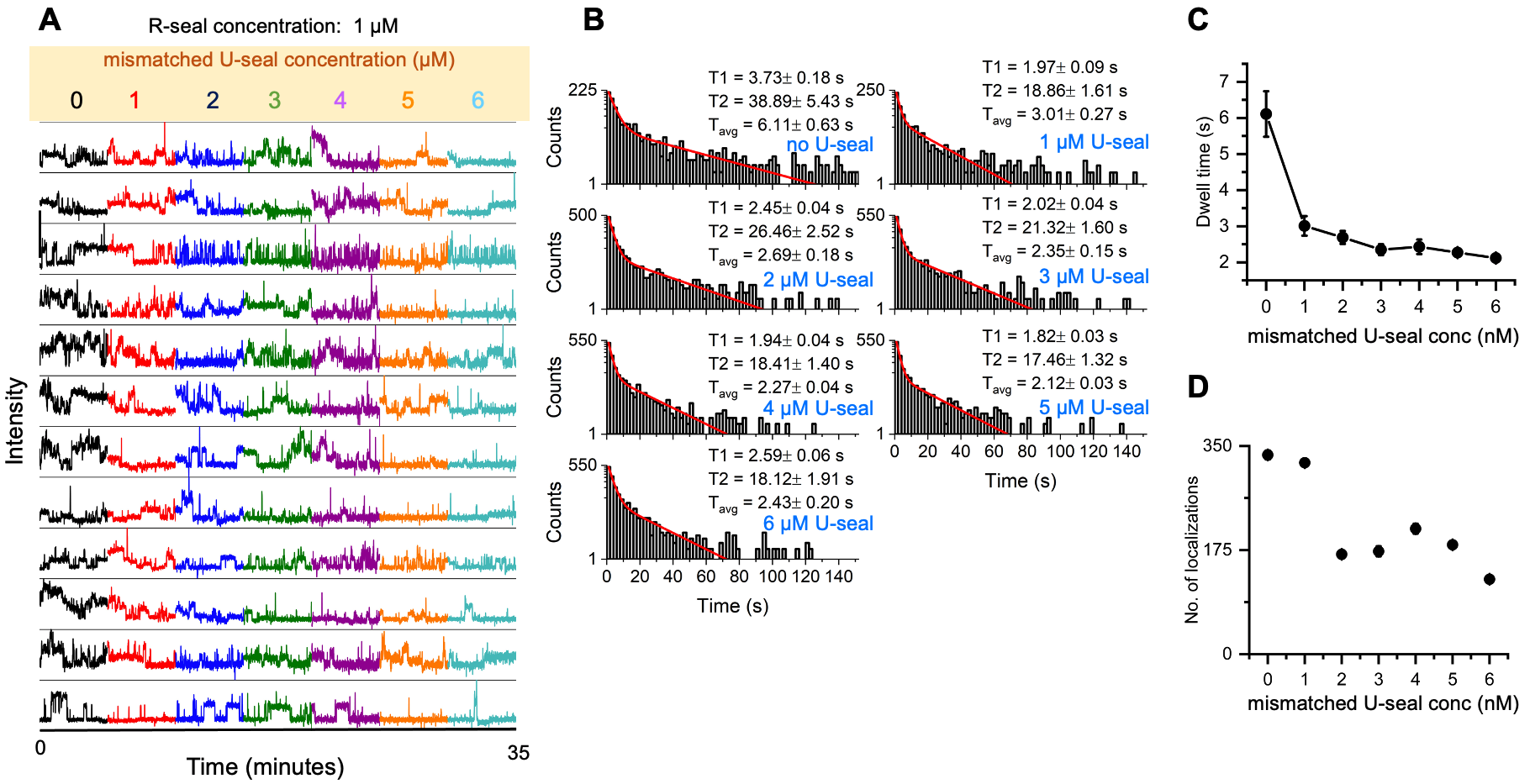

**Fig. S15. Quantifying the effect of competitive inhibition of R-seal binding to a gap due to increasing concentrations of a 1-nt mismatched U-seal: 5-base sequencing.**

**A.** Representative traces for gap-binding of 1 μM of R-seal R13-N^5^N^6^N^7^N^8^N^9^ in presence of 1-5 μM complementary U-seal (U13-T^5^N^6^N^7^N^8^N^9^).

**B.** Dwell time distributions for the different U-seal concentration. Histograms are fitted with double-exponential decay model (red lines).

**C-D.** Mean dwell-times (panel C) and localisation count (panel D) as a function of U-seal concentration.

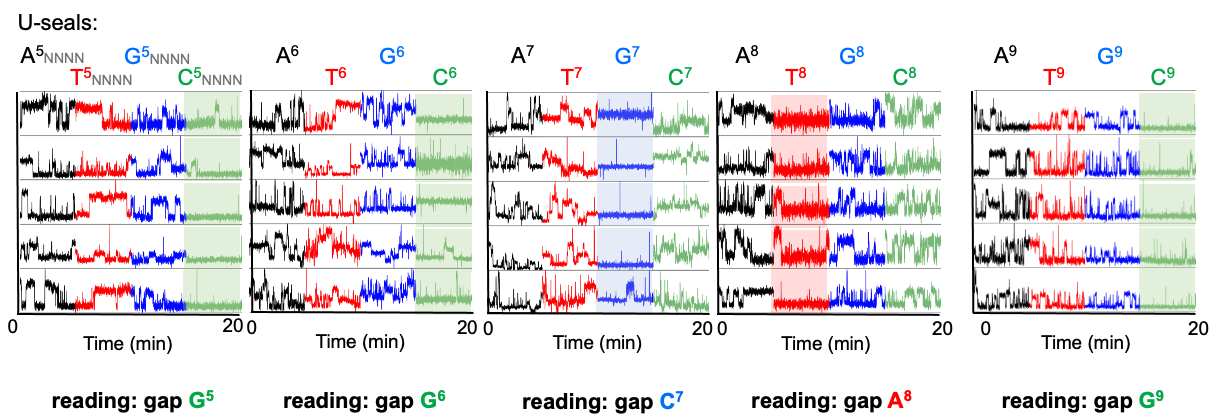

**Fig. S16. Reading five nucleotides using competitive inhibition: additional examples**. Traces for 5-base sequencing are provided for gap positions 5, 6, 7, 8 and 9. Conditions and descriptions as in **Fig. 3H**. In all cases, the R-seal (R13-N^5^N^6^N^7^N^8^N^9^) and the U-seals were kept at 1 μM and 2 μM, respectively. Interrogation with the complementary U-seal led to decrease binding (highlighted traces).

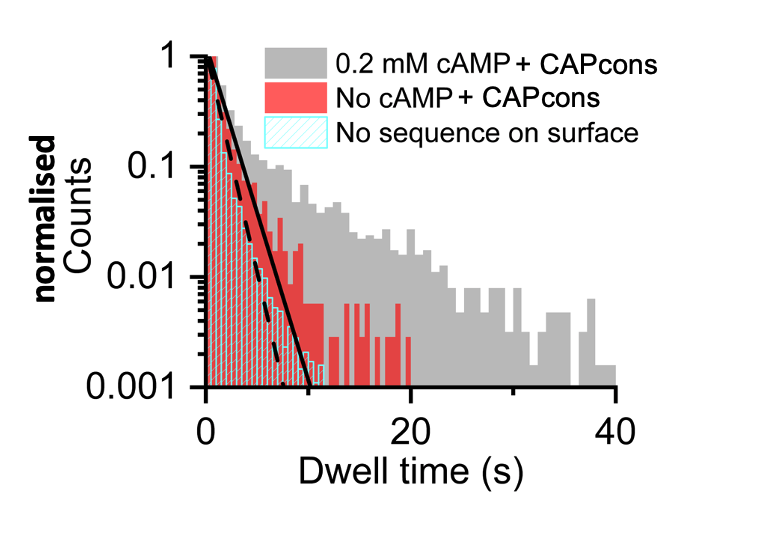

**Fig. S17. Control experiments for specificity of CAP-DNA interactions.** Dwell-time distributions for CAP binding to surface-immobilised molecules in the presence of CAPcons DNA and cAMP (grey columns); in the presence of CAPcons DNA only (no cAMP; pink columns; mean dwell time of 1.47 s); and in the absence of CAPcons DNA and cAMP (cyan hatched bars; mean dwell time of 1.07 s). The comparison of the controls with the sample displaying specific binding supports that assignment of the long lifetime of ~9 sec to specific, cAMP-dependent binding of CAP to its consensus site.

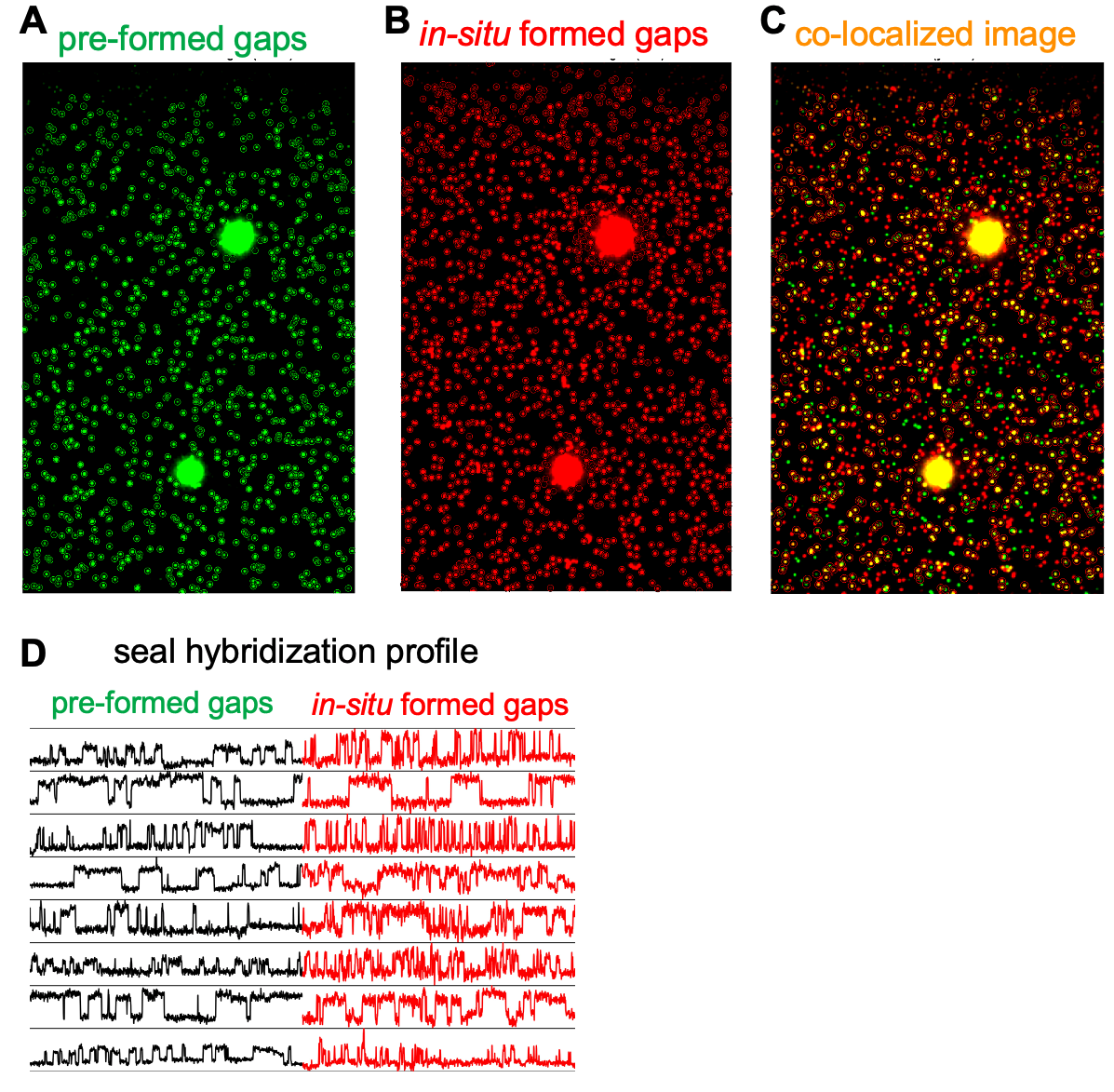

**Fig. S18. Efficiency and functionality of in-situ DNA gap formation.**

**A.** Localisation image of surface-immobilised 8-nt gapped DNA molecules.

**B.** Localisation image of *in-situ* formed surface-immobilised 8-nt gapped DNA molecules (same field-of-view).

**C.** Superimposed images of panels A and B; colocalised molecules appear yellow.

**D.** In black: time-traces of transient seal hybridization to pre-formed surface-immobilised gapped DNA. In red: time-traces of transient seal hybridization to the *in-situ* formed gapped DNA. The binding patterns pre-formed and in-situ formed DNA gaps are similar.

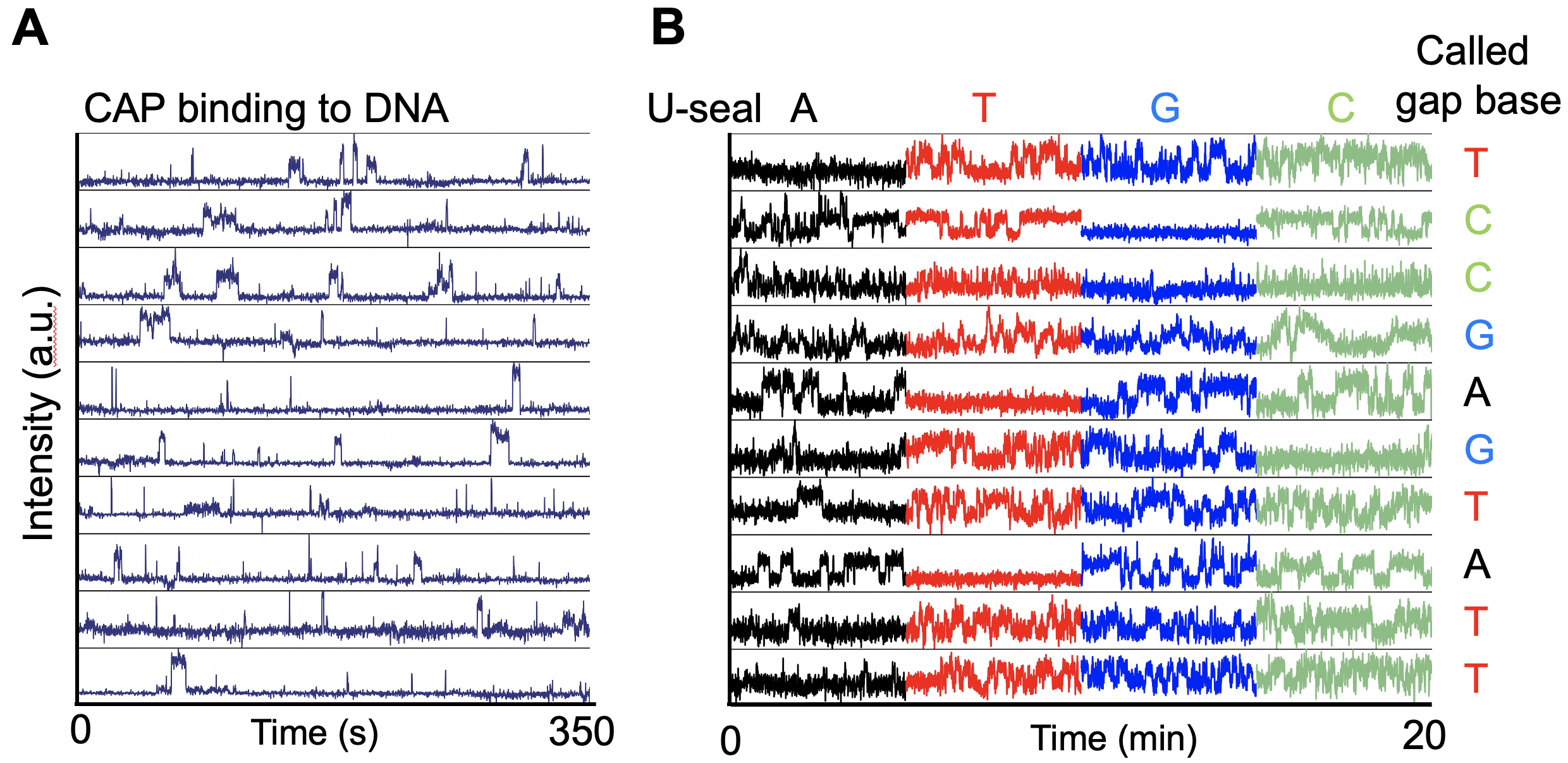

**Fig. S19. Representative traces for CAP-DNA interactions of the library followed by sequencing.**

**A.** Time-traces of CAP binding to members of the library of 4 DNA variants for position 5 within the CAPcons sequence. The assay employed 2.5 nM of Alexa647-labelled CAP.

**B.** Traces obtained from interrogation of the same DNA molecule as shown in the panel A using competitive inhibition of gap binding. The assay used 200 nM of R seal R9-CAP-N^5^-Cy5 and 600 nM of 4 competing U-seals (black, red, blue and green for seals featuring A, T, G or C bases sequences, respectively, at the corresponding sequencing position).

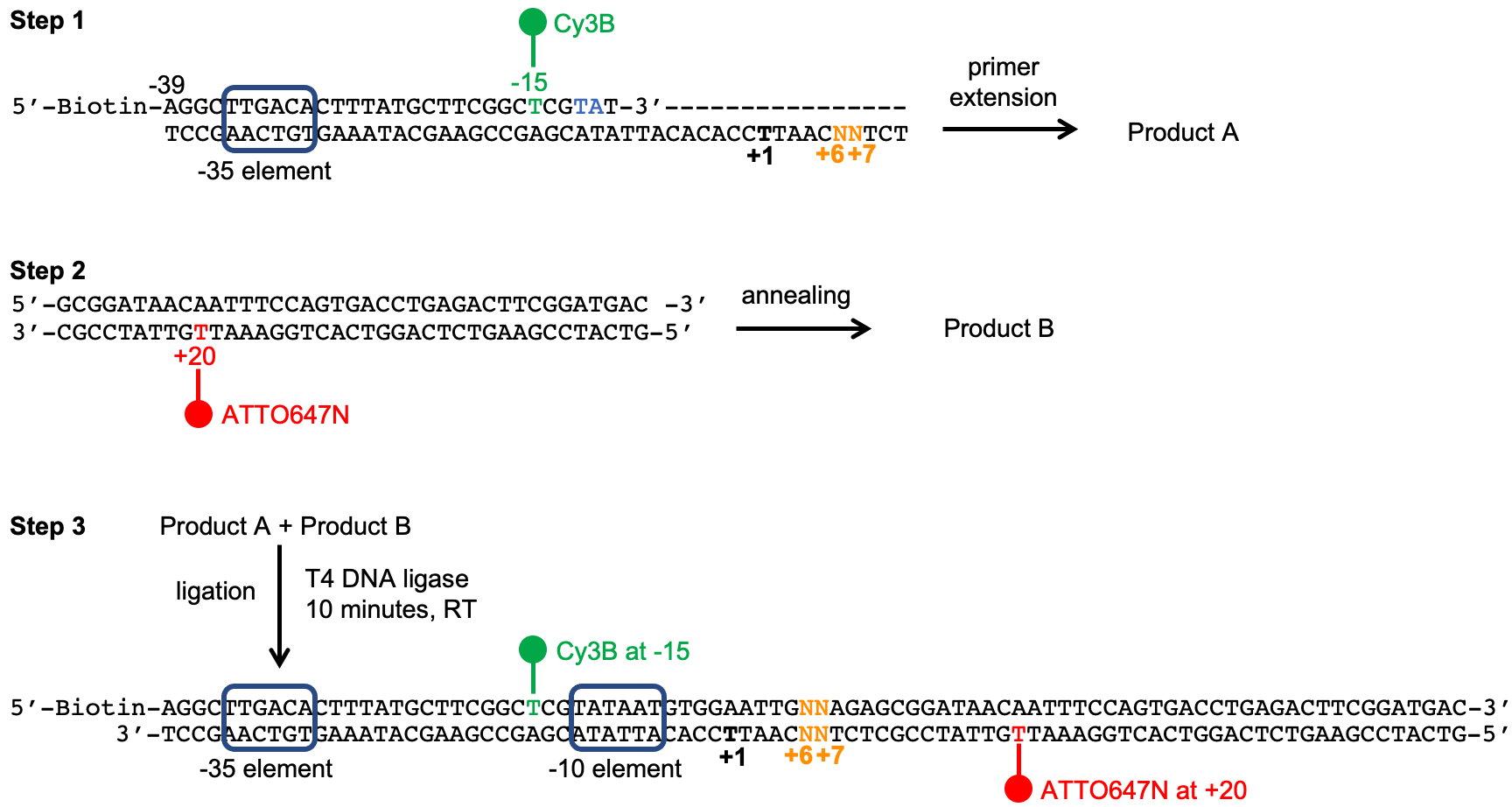

**Fig. S20. Preparation of library of 16 labelled lacCONS promoter DNA variants.** The library was prepared in 3 steps. **Step 1** involves a primer extension of a Cy3B-modified strand annealed on a template strand with degenerate bases at sites corresponding to positions +6 and +7 from the transcription start site; the extended product is then purified. **Step 2** involves preparation of another duplex fragment, wherein the bottom strand is labelled with ATTO647N at the site corresponding to position +20 form the transcription start site. In **Step 3**, the two dsDNA fragments are ligated using a high-performance T4 DNA ligase to form the final doubly labelled promoter dsDNA fragment.

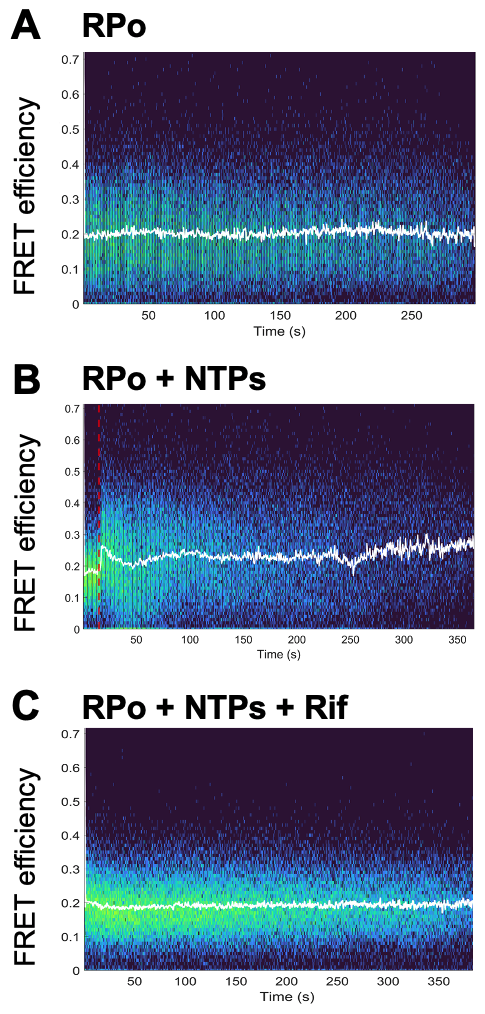

**Fig. S21. Time evolution of FRET efficiencies for transcription complexes at different conditions.** FRET time-traces have been superimposed to produce heat maps that show the time evolution of the FRET signals.

**A.** Open complexes (RPo) prepared using the labelled promoter library and RNAP were immobilized on surface and imaged using alternating-laser excitation to obtain their E_FRET_. The mean E_FRET_ was ~0.22 and does not change during the recording time. N=112.

**B.** Upon adding 500 μΜ ΑpA and 200 μM of all NTPs to the surface-immobilized RPo complexes, a large fraction of the immobilised RPo show an increase in their FRET efficiency at ~15 sec due to the transcription reaction; the following FRET decrease (to a minimum reached in ~50 sec) is due to processes that reduce FRET by populating transcription complexes with E_FRET_ of ~0.2 (mainly abortive RNA release, RNA backtracking and promoter escape; see **Fig. 5A**). The FRET signal also reaches a second local maximum at ~100 sec, likely reflecting a second round of RNA synthesis; beyond that time point, fluctuations are lost due to the desynchronisation of the transcribing complexes. N=159.

**C.** Blocking initial transcription using rifampicin leads to a profile and mean E_FRET_ similar to that for RPo, confirming that the FRET changes observed in the presence of NTPs are due to the transcription reaction. N=117.

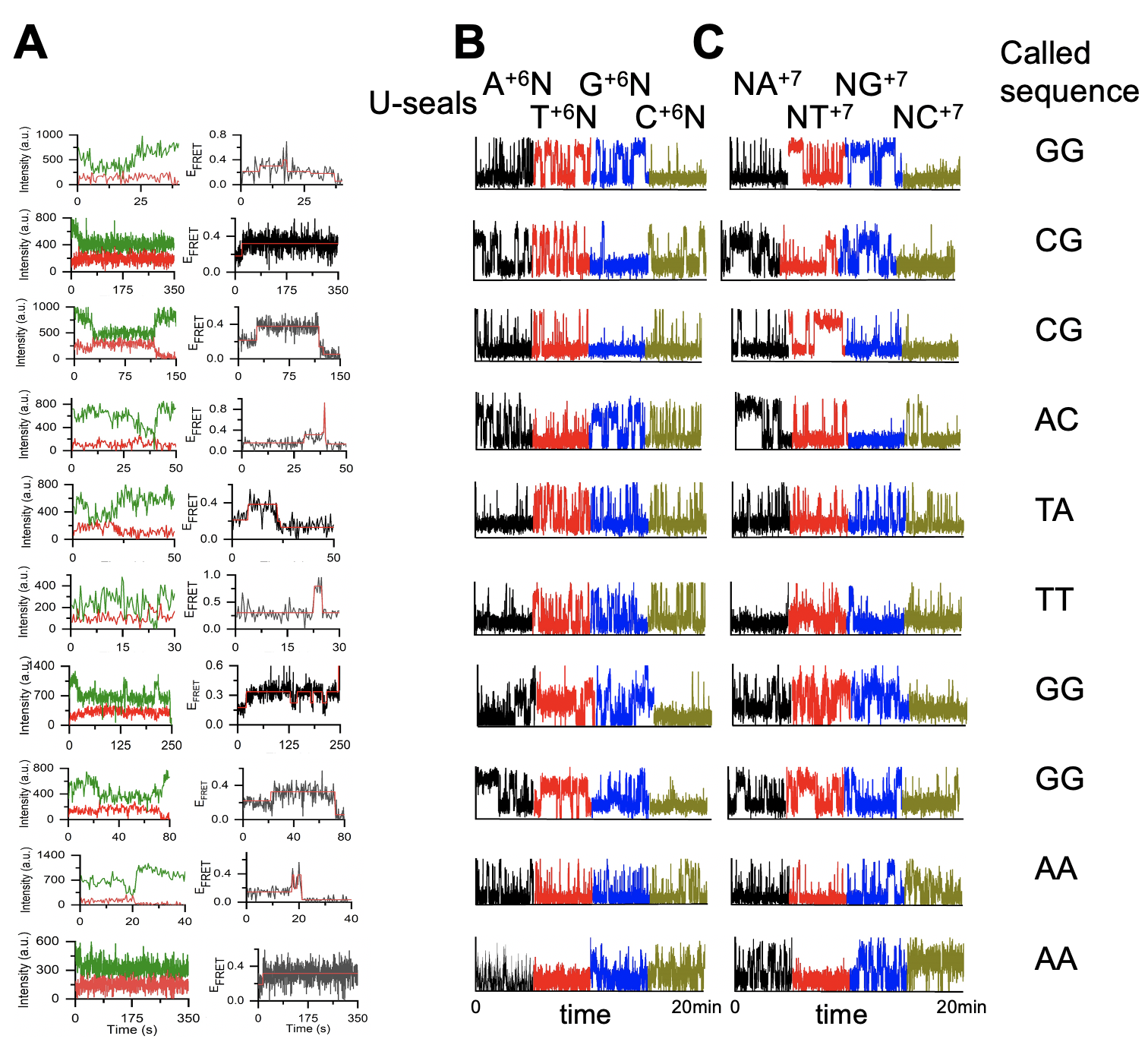

**Fig. S22. Example time-traces obtained from SPIN-seq analysis of the promoter library.**

**A.** **Left,** FRET donor and FRET acceptor intensities upon donor excitation exhibiting anti-correlated changes characteristic of FRET interactions between the two fluorophores. **Right,** FRET efficiency time-traces showing transitions between a low-FRET state (E_FRET_ ~0.2) and an intermediate-FRET state (E_FRET_~0.4).

**B.** Sequencing position +6 using a R-seal (featuring N^+6^N^+7^ bases) and a set of competing U-seals (A^+6^N^+7^, T ^+6^N^+7^, G ^+6^N^+7^, C^+6^N^+7^).

**C.** Similar interrogation at position +7 for the same molecule using the same R-seal and another set of competing U-seals (N^+6^A^+7^, N ^+6^T^+7^, N ^+6^G^+7^, N^+6^C^+7^). In the right panel identified sequence for the same corresponding molecules.

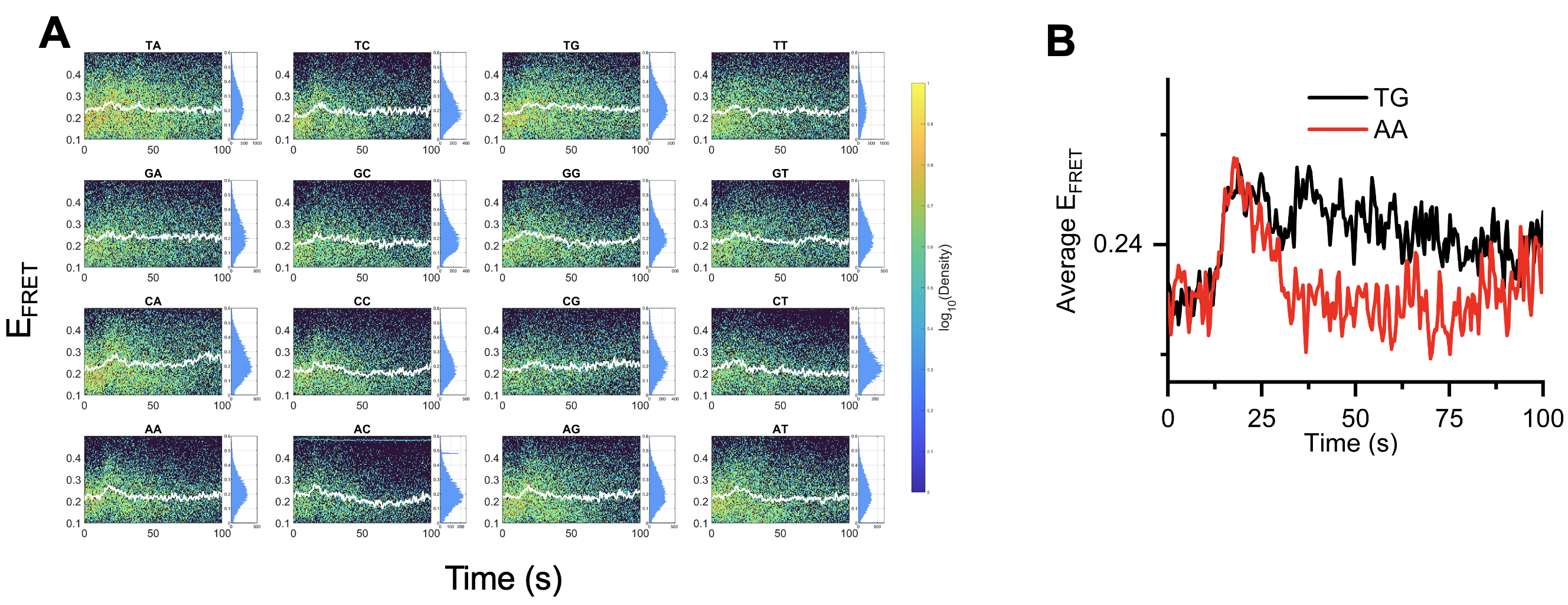

**Fig. S23. Sequence-dependence of FRET changes along the transcription reaction for the lacCONS promoter 16-variant library.**

**A.** FRET timetrace heatmap featuring traces for all transcriptionally active molecules for all 16 sequences. White line: mean FRET as a function of time. Sequences exhibiting frequent and/or long dwells at the paused state with E*~0.4 (e.g., TG and CG) reach a maximum mean value after 15-20 sec and show a very slow decrease from this value thereafter. In contrast, sequences exhibiting short/infrequent pausing (e.g., CT, AA, AC) reach a maximum mean value within 15-20 sec but return to the starting value much more rapidly (in 10-20 sec). Sequences exhibiting elevated propensity for promoter escape (class I) and cycling (class II) behavior show a distinctive feature of rapid FRET increase to the max FRET value, followed by a rapid decrease to the initial FRET value.

**B.** Overlayed average FRET efficiencies across all traces for two sequences, TG (black) and AA (red). Both the sequences exhibit similar temporal profile in terms of reaching high E_FRET_ value upon addition of NTP (~15 sec). Escalated pausing sequences TG retains the high E_FRET_ value throughout the duration. However, AA sequences shows decrease in E_FRET_ after some time.

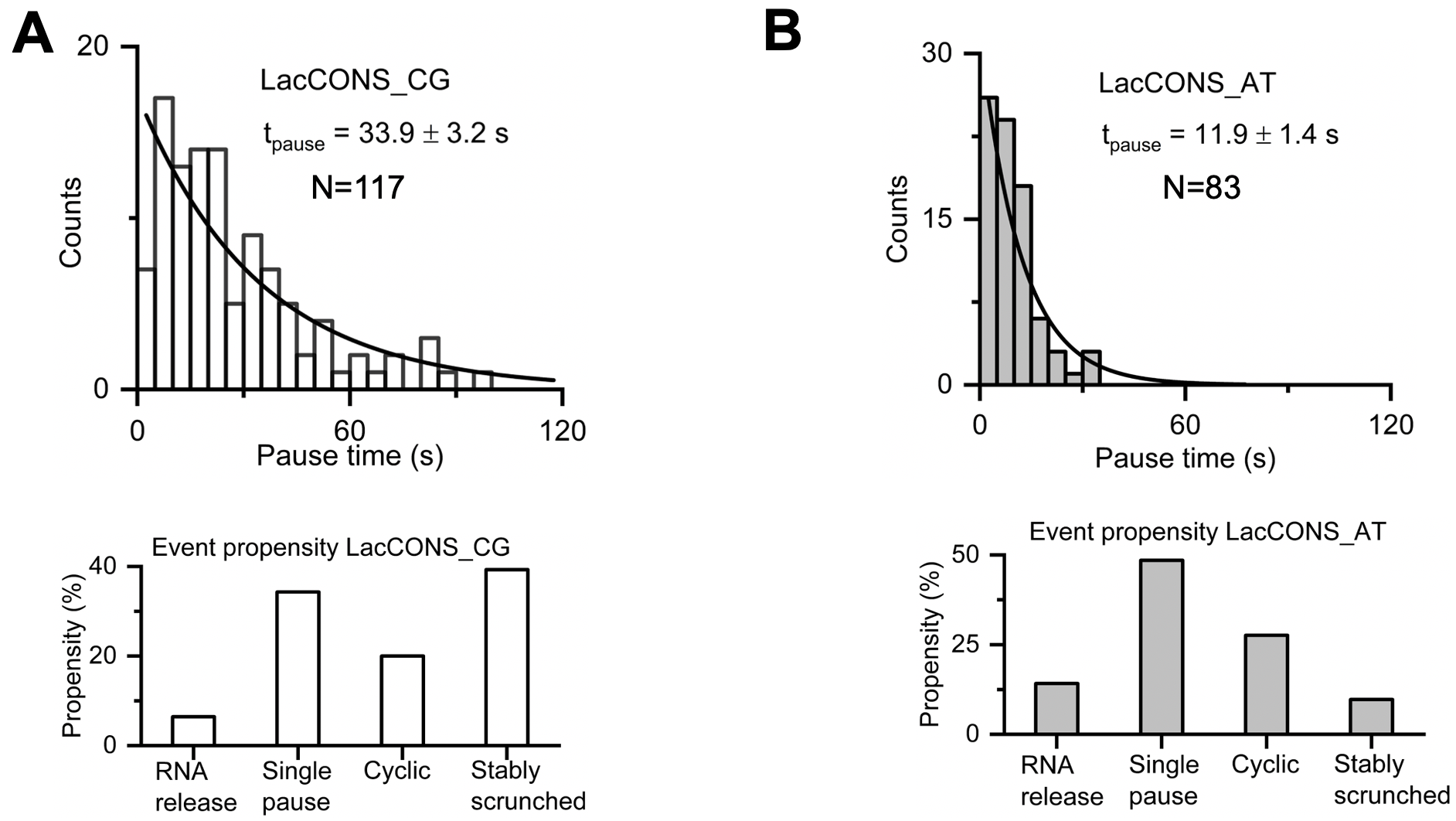

**Fig. S24. Verification of sequence-dependent effects on transcription initiation.** Analysis of two lacCONS promoter variants labelled at -15 and +20 position with Cy3B and ATTO647N, respectively (as for the promoter library). Variants lacCONS-CG and lacCONS-AT were associated with long and short pause duration, respectively, as seen using SPIN-Seq.

**A.** Pause-duration distribution (top) and distribution between classes (bottom) for variant lacCONS-CG (having C and G at positions +6 and +7, respectively). Single-exponential fitting analysis shows of pause lifetime of ~34 s. The variant is associated with pronounced tendency for stable scrunching and single-pause events.

**B.** Pause-duration distribution (top) and distribution between classes (bottom) for variant lacCONS-AT (having A and T at positions +6 and +7, respectively). Single-exponential fitting analysis shows of pause lifetime of ~12 s. The variant is associated with tendency for cyclic events and single-pause events.

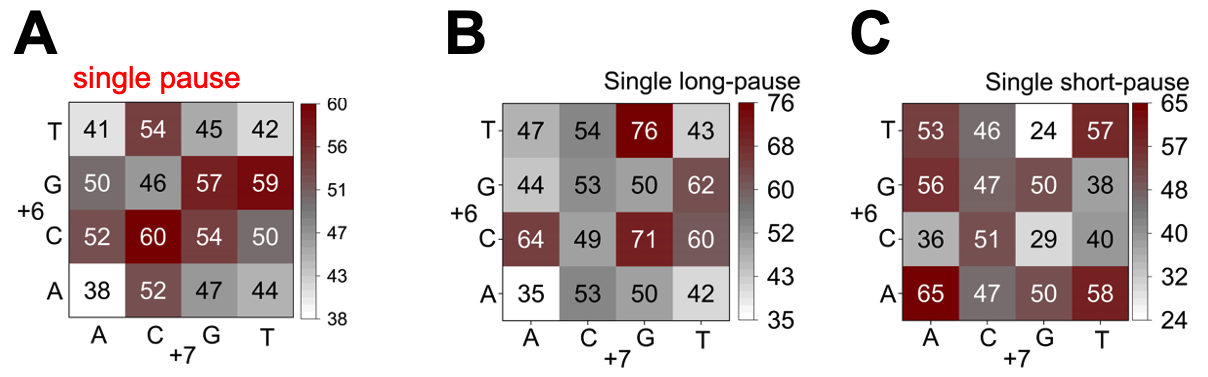

**Fig. S25. Propensity for promoter escape and single-pause events showing pauses of substantially different duration.**

**A.** Propensity of 16 lacCONS promoters to give Class-III events (“single-pause” events).

**B.** Fraction of Class-III events that show a single long-pause (traces showing >10 s single pause duration).

**C.** Propensity of 16 lacCONS promoters to give Class-III events with single short-pause (traces showing 1-10 s single pause duration).

Table S1. DNA sequences used in this work.

| **Experiments/construct** | **Strand Name** | **Sequence** |
| --- | --- | --- |
| Formation of 8-nt gapped DNA (Fig. 2A, 2C, and 2D) | Gap-(1-65)-G^30^-B^Bio,1^ | 5’ CAG TCA AGA GCC TGA GGC AGC AGA TTA TCG ACG GTG GAT GTA TGG TAA TGG GAC GAA GAA TGA GG–biotin 3’ |
|  | Gap-(1-27)-Top^Cy3B, 18^ | 5’ CCT CAT TCT TCG TCC CA**T(Cy3B)** TAC CAT ACA 3’ |
|  | Gap-(36-65)-Top | 5’ CGA TAA TCT GCT GCC TCA GGC TCT TGA CTG 3’ |
| Template for formation of 8-nt gapped DNA with T at position 3 of the gap (Fig. 2C and 2D) | Gap-(1-65)-T^30^-B^Bio,1^ | 5’ CAG TCA AGA GCC TGA GGC AGC AGA TTA TCG ACG GTT GAT GTA TGG TAA TGG GAC GAA GAA TGA GG–biotin 3’ |
| Template for formation of 8-nt gapped DNA with A at position 3 of the gap (Fig. 2C and 2D) | Gap-(1-65)-A^30^-B^Bio,1^ | 5’ CAG TCA AGA GCC TGA GGC AGC AGA TTA TCG ACG GTA GAT GTA TGG TAA TGG GAC GAA GAA TGA GG–biotin 3’ |
| Template for formation of 8-nt gapped DNA with C at position 3 of the gap (Fig. 2C and 2D) | Gap-(1-65)-C^30^-B^Bio,1^ | 5’ CAG TCA AGA GCC TGA GGC AGC AGA TTA TCG ACG GTC GAT GTA TGG TAA TGG GAC GAA GAA TGA GG–biotin 3’ |
| 8 nt fluorescent seals (Fig. 2A, 2C, and 2D) | S8-C^3^ | 5’ATTO647N -TCC ACC GT 3’ |
|  | S8-G^3^ | 5’ATTO647N -TCG ACC GT 3’ |
|  | S8-A^3^ | 5’ATTO647N -TCA ACC GT 3’ |
|  | S8-T^3^ | 5’ATTO647N -TCT ACC GT 3’ |
| 8-nt R-seal for one-base reading (Fig. 3B) | R8-N^3^ | 5’ATTO647N -TCN ACC GT 3’ |
| Set of 4 8-nt U-seals for one-base reading (Fig. 3B) | U8-A^3^ | 5’ TCA ACC GT 3’ |
|  | U8-T^3^ | 5’ TCT ACC GT 3’ |
|  | U8-G^3^ | 5’ TCG ACC GT 3’ |
|  | U8-C^3^ | 5’ TCC ACC GT 3’ |
| Formation of 13-nt gap gapped DNA (Fig. 3E) | Gap-(1-65)-B^Bio,1^ | 5’ CAG TCA AGA GCC TGA GGC AGC AGA TTA TCG ACG GTG GAT GTA TGG TAA TGG GAC GAA GAA TGA GG–biotin 3’ |
|  | Gap-(1-27)-Top^Cy3B, 25^ | 5’ CCT CAT TCT TCG TCC CA**T(Cy3B)** TAC CAT ACA 3’ |
|  | Gap-(41-65)-Top | 5’ A TCT GCT GCC TCA GGC TCT TGA CTG 3’ |
| 13-nt R-seal for 3-base reading (Fig. 3E) | R13-N^5^N^6^N^7^ | 5’ - Atto647N **-**TCC ANN NTC GAT A- BHQ1 - 3’ |
| Set of 12 13-nt U-seals for 3-base reading (Fig. 3E) | U13-A^5^N^6^N^7^ | 5’ TCC AAN NTC GAT A 3’ |
|  | U13-T^5^N^6^N^7^ | 5’ TCC ATN NTC GAT A 3’ |
|  | U13-G^5^N^6^N^7^ | 5’ TCC AGN NTC GAT A 3’ |
|  | U13-C^5^N^6^N^7^ | 5’ TCC ACN NTC GAT A 3’ |
|  | U13-N^5^A^6^N^7^ | 5’ TCC ANA NTC GAT A 3’ |
|  | U13-N^5^T^6^N^7^ | 5’ TCC ANT NTC GAT A 3’ |
|  | U13-N^5^G^6^N^7^ | 5’ TCC ANG NTC GAT A 3’ |
|  | U13-N^5^C^6^N^7^ | 5’ TCC ANC NTC GAT A 3’ |
|  | U13-N^5^N^6^A^7^ | 5’ TCC ANN ATC GAT A 3’ |
|  | U13-N^5^N^6^T^7^ | 5’ TCC ANN TTC GAT A 3’ |
|  | U13-N^5^N^6^G^7^ | 5’ TCC ANN GTC GAT A 3’ |
|  | U13-N^5^N^6^C^7^ | 5’ TCC ANN CTC GAT A 3’ |
| 13-nt R-seal for 5-base reading (Fig. 3G) | R13-N^5^N^6^N^7^N^8^N^9^ | 5’- ATTO647N **-**TCC ANN NNN GAT A- BHQ1 - 3’ |
| Set of 20 13-nt U-seals for 5-base reading (Fig. 3G) | U13-A^5^N^6^N^7^N^8^N^9^ | 5’ TCC AAN NNN GAT A 3’ |
|  | U13-T^5^N^6^N^7^N^8^N^9^ | 5’ TCC ATN NNN GAT A 3’ |
|  | U13-G^5^N^6^N^7^N^8^N^9^ | 5’ TCC AGN NNN GAT A 3’ |
|  | U13-C^5^N^6^N^7^N^8^N^9^ | 5’ TCC ACN NNN GAT A 3’ |
|  | U13-N^5^A^6^N^7^N^8^N^9^ | 5’ TCC ANA NNN GAT A 3’ |
|  | U13-N^5^T^6^N^7^N^8^N^9^ | 5’ TCC ANT NNN GAT A 3’ |
|  | U13-N^5^G^6^N^7^N^8^N^9^ | 5’ TCC ANG NNN GAT A 3’ |
|  | U13-N^5^C^6^N^7^N^8^N^9^ | 5’ TCC ANC NNN GAT A 3’ |
|  | U13-N^5^N^6^A^7^N^8^N^9^ | 5’ TCC ANN ANN GAT A 3’ |
|  | U13-N^5^N^6^T^7^N^8^N^9^ | 5’ TCC ANN TNN GAT A 3’ |
|  | U13-N^5^N^6^G^7^N^8^N^9^ | 5’ TCC ANN GNN GAT A 3’ |
|  | U13-N^5^N^6^C^7^N^8^N^9^ | 5’ TCC ANN CNN GAT A 3’ |
|  | U13-N^5^N^6^N^7^A^8^N^9^ | 5’ TCC ANN NAN GAT A 3’ |
|  | U13-N^5^N^6^N^7^T^8^N^9^ | 5’ TCC ANN NTN GAT A 3’ |
|  | U13-N^5^N^6^N^7^G^8^N^9^ | 5’ TCC ANN NGN GAT A 3’ |
|  | U13-N^5^N^6^N^7^C^8^N^9^ | 5’ TCC ANN NCN GAT A 3’ |
|  | U13-N^5^N^6^N^7^N^8^A^9^ | 5’ TCC ANN NNA GAT A 3’ |
|  | U13-N^5^N^6^N^7^N^8^T^9^ | 5’ TCC ANN NNT GAT A 3’ |
|  | U13-N^5^N^6^N^7^N^8^G^9^ | 5’ TCC ANN NNG GAT A 3’ |
|  | U13-N^5^N^6^N^7^N^8^C^9^ | 5’ TCC ANN NNC GAT A 3’ |
| DNAs for annealing of CAPcons dsDNA (Fig. 4D) | CAPcons-(1-49)-C^25^-B^Bio,49^ | 5’ biotin-GTG CCT AAA ATG TGA TCT AGA TCA CAT TTA TTG CGT AGA GCT CAC TGC C 3’ |
|  | CAPcons-(1-49)- G^25^-Top^Cy3B,13^ | 5’ GGC AGT GAG CTC **T(Cy3B)** AC GCA ATA AAT GTG ATC TAG ATC ACA TTT TAG GCA C 3’ |
| DNAs for annealing of CAPcons dsDNA G:C variant (Fig. 4E) | CAPcons-(1-49)-G^25^-B^Bio,49^ | 5’ biotin-GTG CCT AAA ATG TGA TCT AGA TCA GAT TTA TTG CGT AGA GCT CAC TGC C 3’ |
|  | CAPcons-(1-49)-C^25^-Top^Cy3B,13^ | 5' GGC AGT GAG CTC **T(Cy3B)** AC GCA ATA AAT CTG ATC TAG ATC ACA TTT TAG GCA C 3’ |
| DNA for preparation of CAP variant library at position 5 of the CAP consensus (Fig. 4F) | CAPcons-(1-49)-N^25^-B^Bio,49^ | 5’ biotin-GTG CCT AAA ATG TGA TCT AGA TCA NAT TTA TTG CGT AGA GCT CAC TGC C 3’ |
|  | CAPcons-(1-20)-Top^Cy3B,11^ | 5’ GGC AGT GAG C**TCy3B)** C TAC GCA AT 3’ |
| Flanking strands for *in situ* gap formation on CAPcons library (Fig. 4F) | CAPcons-(30-49)-Top | 5’ CTA GAT CAC ATT TTA GGC AC 3’ |
|  | CAPcons-(1-20)-Top^Cy3B,11^ | 5’ GGC AGT GAG C**T(Cy3B)** C TAC GCA AT 3’ |
| 9-nt CAPcons R-Seal (Fig. 4H) | R9-CAP-N^5^ | 5’ Cy5 - AAA TNT GAT 3’ |
| Set of 4 9-nt CAPcons U-seals (Fig. 4H) | U9-CAP-A^5^ | 5’ AAA TAT GAT 3’ |
|  | U9-CAP-T^5^ | 5’ AAA TTT GAT 3’ |
|  | U9-CAP-G^5^ | 5’ AAA TGT GAT 3’ |
|  | U9-CAP-C^5^ | 5’ AAA TCT GAT 3’ |
| DNAs for generation of *lac*CONS variant promoter (Fig. 5B) | lacCONS(-39/-10)-Top^Cy3B,-15; Bio,-39^ | 5’ biotin- AGG CTT GAC ACT TTA TGC TTC GGC **T(Cy3B)** CG TAT 3’ |
|  | lacCONS(-39/+10)-N^+6^N^+7^-B | 5’ TCT NNC AAT TCC AAC ACG GCT ACG AGC CGA AGC ATA AAG TG 3’ |
|  | lacCONS(+11/+49)-B^Atto647N,+20^ | 5’ GTC ATC CGA AGT CTC AGG TCA CTG GAA AT**T(ATTO647N)** GTT ATC CGC 3’ |
|  | lacCONS(+11/+49)-Top | 5’ GCG GAT AAC AAT TTC CAG TGA CCT GAG ACT TCG GAT GAC 3’ |
| Flanking strands for lacCONS gap formation around positions +6/+7 (Fig. 5C) | lacCONS(+11/+49)-B | 5’ GTC ATC CGA AGT CTC AGG TCA CTG GAA ATT GTT ATC CGC 3’ |
|  | lacCONS(-39/+2)-B | 5’ TTC CAC ACA TTA TAC GAG CCG AAG CAT AAA GTG TCA AGC CT 3’ |
| 8-nt lacCONS R-seal (Fig. 5C) | R8-lacCONS(+3/+9)-N^+7^N^+6^-B | 5’ ATTO647N-TCT NNC AA 3’ |
| Set of 8 8-nt lacCONS U-seals (Fig. 5C) | U8-lacCONS(+3/+9) -A^+7^-N^+6^-B | 5’ TCT ANC AA 3’ |
|  | U8-lacCONS(+3/+9) -T^+7^-N^+6^-B | 5’ TCT TNC AA 3’ |
|  | U8-lacCONS(+3/+9) -G^+7^-N^+6^-B | 5’ TCT GNC AA 3’ |
|  | U8-lacCONS(+3/+9) -C^+7^-N^+6^-B | 5’ TCT CNC AA 3’ |
|  | U8-lacCONS(+3/+9) -N^+7^-A^+6^-B | 5’ TCT NAC AA 3’ |
|  | U8-lacCONS(+3/+9) -N^+7^-T^+6^-B | 5’ TCT NTC AA 3’ |
|  | U8-lacCONS(+3/+9) -N^+7^-G^+6^-B | 5’ TCT NGC AA 3’ |
|  | U8-lacCONS(+3/+9) -N^+7^-C^+6^-B | 5’ TCT NCC AA 3’ |
| DNAs for preparation of *lac*CONS A^+6^T^+7^ variant (fig. S24) | lacCONS(-39/+10)-A^+6^T^+7^-B^Cy3B,-15; Bio,-39^ | 5’ Biotin-AGG CTT GAC ACT TTA TGC TTC GGC **T(Cy3B)** CG TAT AAT GTG TGG AAT TGA TAG A 3’ |
|  | lacCONS(-39/+10)-A^+6^T^+7^-Top | 5’ TCT ATC AAT TCC ACA CAT TAT ACG AGC CGA AGC ATA AAG TGT CAA GCC T 3’ |
| DNAs for preparation of *lac*CONS C^+6^G^+7^ variant (fig. S24) | lacCONS(-39/+10)-C^+6^G^+7^- B^Cy3B,-15; Bio,-39^ | 5’ Biotin - AGG CTT GAC ACT TTA TGC TTC GGC **T(Cy3B)** CG TAT AAT GTG TGG AAT TGC GAG A 3’ |
|  | lacCONS(-39/+10)-C^+6^G^+7^-Top | 5’ TCT CGC AAT TCC ACA CAT TAT ACG AGC CGA AGC ATA AAG TGT CAA GCC T 3’ |

Underlined sequences denote the part of a long strand that form the ssDNA gap in a gapped DNA. Bold typeface denotes a labelled position along with the fluorophore attached (the fluorophores used is in a parenthesis). N indicates a degenerate base. Top: top strand. B: bottom strand. Bio: biotin. Underlined sequence: template sequence for the DNA gap formed by the given template strand.

Table S2. Accuracies for base-calling for the competitive-inhibition-based assay

| **Accuracy for single-base sequencing using competitive-inhibition assay** | | |
| --- | --- | --- |
| **Gap Base** | **Accuracy (%), individual gaps** | **Accuracy (%) mixed gaps** |
| A | 94.2 (274/291) | 97.9 (184/188) |
| T | 97.2 (383/394) | 98.8 (243/246) |
| G | 99.6 (500/502) | 99.6 (224/225) |
| C | 96.6 (736/762) | 96.9 (190/196) |
| All bases | 97.1 (1893/1949) | 98.5 (841/855) |
| **Accuracy for 3-base sequencing using competitive-inhibition assay** | | |
| **Gap Base** | **Accuracy (%)** | |
| G^5^ | 96.4 (240/249) | |
| G^6^ | 98.1 (256/261) | |
| C^7^ | 95.0 (489/515) | |
| All bases | 96.5 (765/793) | |
| **Accuracy for 5-base sequencing using competitive binding assay** | | |
| **Gap Base** | **Accuracy (%)** | |
| G^5^ | 95.1 (195/205) | |
| G^6^ | 97.9 (464/474) | |
| C^7^ | 97.1 (438/451) | |
| A^8^ | 94.9 (352/371) | |
| G^9^ | 97.1 (233/240) | |
| All bases | 96.6 (1682/1741) | |

Table S3. Kinetic analysis of CAP-DNA interactions based on single-molecule binding time-traces.

| **Kinetic analysis for CAP binding to consensus and single mutant DNA sequences** | | | |
| --- | --- | --- | --- |
| **Sequence** | **Dwell times t_1_ and t_2_ (s)** | **On-rate (M^-1^s^-1^)** | **K_d_ (nM)** |
| CAPcons_G | 0.55 ± 0.31  6.42 ± 0.35 | 6.8 x 10^6^ | 20.2 ± 1.3 |
| CAPcons_C (consensus) | 1.75 ± 0.42  8.71 ± 0.50 | 1.3 x 10^7^ | 9.1 ± 0.6 |
| **Kinetic analysis for CAP-binding to the library of CAPcons DNA variants** | | | |
| **Sequence** | **Dwell times t_1_ and t_2_ (s)** | **On-rate (M^-1^s^-1^)** | **K_d_ (nM)** |
| CAPcons_A | 0.94 ± 0.03  4.39 ± 0.52 | 8.32 x 10^6^ | 28.9 ± 5.4 |
| CAPcons_T | 0.79 ± 0.07  5.42 ± 1.45 | 7.28 x 10^6^ | 25.4 ± 6.3 |
| CAPcons_G | 0.85 ± 0.05  5.82 ± 1.31 | 8.24 x 10^6^ | 21.2 ± 6.1 |
| CAPcons_C  (consensus) | 0.94 ± 0.01  7.91 ± 0.57 | 1.2 x 10^7^ | 11.2 ± 2.2 |

**List of supplemental references**
